# Three new species of nematodes from the syconia of *Ficus racemosa* in southern India

**DOI:** 10.1101/2021.05.17.444399

**Authors:** Satyajeet Gupta, Qudsia Tahseen, Renee M. Borges

**Affiliations:** Centre for Ecological Sciences, Indian Institute of Science, Bangalore 560012, India; Department of Zoology, Aligarh Muslim University, Aligarh 202002, India

**Keywords:** Fig nematodes, fig wasp, morphology, new species, morphometrics, molecular characterization

## Abstract

*Ficus racemosa* with an Indo-Australasian distribution, has so far been recorded to harbour in its fruits, nematode species of the aphelenchoid genera *Schistonchus*, *Ficophagus* and *Martininema*, and species of diplogastrid genera *Teratodiplogaster* and *Pristionchus*. The Indian species reported so far from *Ficus racemosa* lack comprehensive details on morphology and molecular characterization. In this paper, we describe three new species of nematodes obtained from syconia (enclosed globular infructescence or fruit) of *Ficus racemosa* found in southern India. *Ficophagus glomerata* n. sp. is characterised by small body having b=5.2-9.6, c= 18-23; slightly set-off lip region having well developed cephalic framework; secretory-excretory pore opens near the head; slender stylet with small, rounded/ sloping knobs; ovoid median bulb with relatively posteriorly-placed valve plates; males with sickle-shaped spicules having spatulate or hammer-shaped capitulum, represented by an elongate-ovoid condylus and long digitate rostrum and tail conoid with fine, hair-like terminal spike. *Teratodiplogaster glomerata* n. sp. is characterised by long tubular and narrow stoma with fractal pieces in prostegostom; long rectangular metacorpus; female reproductive system with conspicuous spermatheca and amoeboid sperms; males having short, arcuate spicules and keeled gubernaculum; genital papillae in the configuration of P1, P2, C, P3, P4, P5d, (P6d, P7), P8d, Ph and tail conoid with a terminal or subterminal mucro. *Pristionchus glomerata* n. sp. is characterised by four morphotypes mainly with variations in lip region, stoma, spicules, gubernaculum, and the position of genital papillae. Phylogenetic analyses based upon near-full-length small subunit (SSU) and D2–D3 expansion segments of large subunit (LSU) rRNA genes confirmed affinities with sister species of corresponding genera.

---

Nematodes form the most abundant and diverse metazoans in the world and are associated with meso- and macro-faunal arthropods in various associations, ranging from parasitic to mutualistic (Giblin-Davis *et al*., 2013). These associations have been proposed to lead to speciation and cause increase in nematode diversity and phenotypic plasticity by forming isolated systems (Price, 1980). One such example where such diversification has been observed is the fig–fig wasp–nematode system.

The fig–fig wasp system represents a ∼75 million-year old obligate relationship (Cruaud *et al*., 2012) with over 800 known fig species, each associated with a specific pollinator wasp species, although host-switches and pollinator host sharing has been observed in neotropics (Weiblen, 2002; Machado *et al*., 2005). The enclosed fig inflorescence, called a syconium, forms a microcosm containing different species of wasps, bacteria, fungi, mites, and nematodes (Herre *et al*., 2008). The pollinator wasps are known to be associated with the nematode community comprising of individuals of a single or multiple taxa. The nematodes reported so far from syconia belong to the families, Rhabditidae, Diplogastridae and Aphelenchoididae (Giblin-Davis *et al*., 2006; Gulcu *et al*., 2008; Powers *et al*., 2009; Susoy *et al*., 2016; Kanzaki *et al*., 2018) with associations ranging from phoresy to parasitism. Fig nematodes use fig wasps as their vectors in order to move from one fig (microcosm) to another (Krishnan *et al*., 2010). The nematodes may be transported in the cavity, in intersegmental folds or may cling on to the body surface of the female fig wasp (Giblin-Davis *et al*., 1995). *Parasitodiplogaster*, *Teratodiplogaster*, *Pristionchus*, *Acrostichus*, *Rhabditolaimus* (Diplogastridae); *Bursaphelenchus* (Aphelenchoididae); *Caenorhabditis* (Rhabditidae); *Schistonchus* (Aphelenchoididae); *Ficophagus* (Aphelenchoididae) and *Martininema* (Aphelenchoididae) are the genera known to be phoretic on pollinator fig wasps.

*Ficus racemosa*, the model system in this study, is a monoecious plant species whose syconia, harbour a community of nematodes. The nematode species reported from Indian *F. racemosa* include *Schistonchus racemosa* (Reddy & Rao, 1986), *S. osmani* (Anand, 2002), *S. cuculloracemosus* (Bajaj & Tomar, 2014), *S. flagelloracemosus* (Bajaj & Tomar, 2014), *S. mucroracemosus* (Bajaj & Tomar, 2014), *Teratodiplogaster racemosus* (Bajaj & Tomar, 2015) and *Canalodiplogaster racemosus* (Bajaj & Tomar, 2015); while the species reported from Australian *F. racemosa* include *Ficophagus altermacrophylla* (Kanzaki *et al*., 2010) Davies & Bartholomaeus, 2015; *F. aculeata* (Kanzaki *et al*., 2010) Davies & Bartholomaeus, 2015, *F. fleckeri* (Kanzaki *et al*., 2010) Davies & Bartholomaeus, 2015 and *Teratodiplogaster fignewmani* (Kanzaki, 2009). The diplogastrid species *Pristionchus racemosa* (Susoy *et al*., 2016) was described from syconia of *F. racemosa* from Vietnam.

The species reported from India lack critical morphological data for diagnosis and differentiation (Davies *et al*., 2015). All of them also lack molecular characterization which makes future work with re-isolated lines difficult. Thus, there is a need to describe/re-describe the Indian species of nematodes associated with fig synconia with modern technologies. The present study deals with the morphological characterization of the nematode species isolated from *Ficus racemosa* located in southern India along with their molecular characterization to assess their phylogenetic status.

## Materials and Methods

### Sample collection

The fig syconia in *Ficus racemosa* pass through five developmental phases: phase A (pre-pollination stage, when syconia are at floral bud stage), phase B (pollen-receptive stage, marked by entry of pollinators through ostiole, a tiny opening in the syconium), phase C (development phase of seeds, nematodes and wasps), phase D (wasp dispersal through the opening generated by the male pollinators) and phase E (fruit ripening stage) (Ranganathan *et al*., 2010). Nematodes were collected from the mid-C phase syconia of *F. racemosa* trees in November 2015 from the campus of the Indian Institute of Science, Bangalore, Karnataka, India at coordinates 13.0219° N, 77.5671° E.

### Isolation and observation

Syconia were washed and cut into small pieces using a scalpel. The latex produced by the middle layer of the syconium was absorbed using a tissue paper and the pieces of syconia were immersed in sterilized water for 15–20 min in a Petri dish. Nematodes were hand-picked under a stereoscopic microscope in sterilized water and later fixed in 4% formalin. The fixed nematodes were then dehydrated using glycerol-alcohol (a mixture of 95 parts of 30% ethanol + 5 parts glycerol) solution (Seinhorst *et al*., 1959). Nematodes were mounted permanently on slides using wax ring technique (De Maeseneer & D’ Herde, 1963) for observing under light microscope and conducting confocal imaging using Airyscan LSM 880, Ziess. Drawings were made by tracing the confocal images while measurements were done using Image J 1.46r.

### Scanning Electron Microscopy (SEM)

SEM imaging was done to elucidate the surface features of the nematodes. The nematodes were first fixed in 2% glutaraldehyde for 24 h and then post-fixed in 2% osmium tetroxide for 2 h in the dark in a refrigerator. The nematodes were then washed with PBS and transferred into BEEM^©^ capsules fitted with nylon mesh of 20 μm pore size to hold the specimens (Bozzola & Russell, 1999). The dehydration was carried out by using a serial gradation of alcohol up to 100% and further the nematodes were critical point dried using CO_2_. Later, the nematodes were mounted on double-sided carbon tape placed on the stub, sputtered with 10 nm gold for 38 s and viewed under SEM JEOL (JSM6510L) at 15 kV at the USIF, Aligarh Muslim University (Aligarh).

### Molecular profiles and phylogeny

Nematode DNA was isolated by picking up a single nematode in 5 µl of worm lysis buffer (50 mM KCl, 10 mM Tris HCL pH 8.3, 2.5 mM MgCl_2_, 0.45% NP-40 and 0.45% Tween-20) as per the protocol of Williams *et al*. (1992) and keeping it in a water bath at 65°C for 45 min followed by 95°C for 15 min. The extracted DNA was stored at −20°C and used as template. Primers (Sigma Aldrich) used to amplify D2/D3LSU segment of all three species, were D2A (ACAAGTACCGTGGGGAAAGTTG) and D2B (TCGGAAGGAACCAGCTACTA) (Kanzaki *et al*., 2009). Primers used to amplify SSU segments for *F. glomerata* n. sp. were SNF (TGGATAACTGTGGTAATTCTAGAGC) (Zeng *et al*., 2007) and SNR (TTACGACTTTTGCCCGGTTC); for *T. glomerata* n. sp. and *P. glomerata* n. sp., these primers were SSUF07 (AAAGATTAAGCCATGCATG) and SSU26R (CATTCTTGGCAAATGCTTCG) (Hermann *et al*., 2006, 2007). The PCR products were sequenced, and the sequences were deposited in the GenBank database (Accession numbers in form of SSU/D2-D3 LSU for *F. glomerata* n. sp. MT903999/ MT903998, for *T. glomerata* n. sp. MT904001/ MT904002 and for *P. glomerata* n. sp. MT904000/ MT903997). The sequences were aligned by ClustalW and Muscle in MEGA 7 software and were further compared with those of other nematode species available at the GenBank database using the BLAST homology search program (Zhao *et al*., 2015 for *Ficophagus* spp.; Kanzaki *et al*., 2014 for *Teratodiplogaster* spp. and Susoy *et al*., 2016 for *Pristionchus* spp). The best model used to generate the Bayesian trees was inferred by using Partition finder ver. 2.1.1. The Akaike-supported model, log likelihood (lnL), Akaike information criterion (AIC), the proportion of invariable sites, and the gamma distribution shape parameters and substitution rates were also used in the phylogenetic analyses. The inferred model was further used to generate phylogenetic trees using Mr. Bayes software ver. 3.2.6 (Huelsenbeck & Ronquist, 2001) by running the chain for 1,000,000 generations and the ‘burnin’ was set at 1,000. We used the Markov Chain Monte Carlo (MCMC) method within a Bayesian framework to estimate the posterior probabilities of the phylogenetic trees (Larget & Simon, 1999) using 50% majority rule; the trees were visualized using Fig Tree ver. 1.4.3 (Rambaut, 2006).

## Results

All new species described in this paper have been assigned the specific name “*glomerata*” since *Ficus glomerata* is a synonym of *Ficus racemosa*.

***Ficophagus glomerata* n. sp.** (Figs. 1-5)

**Fig. 1.**
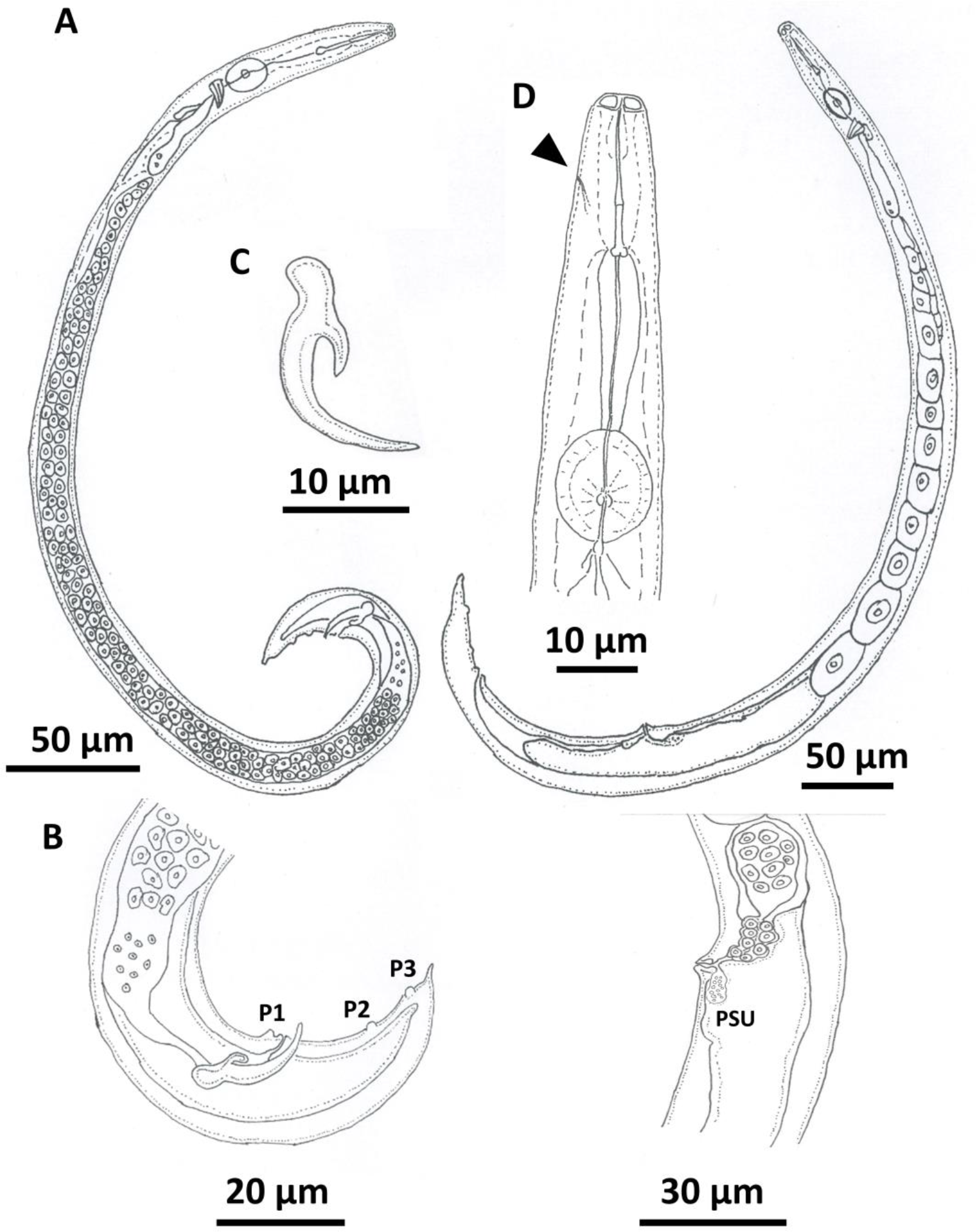
*Ficophagus glomerata* n. sp. lateral view Male A: Habitus, B: Tail region, C: Spicule, D: Anterior region (similar in case of female); Female E: Habitus and F: Vulval region.

**Fig. 2.**
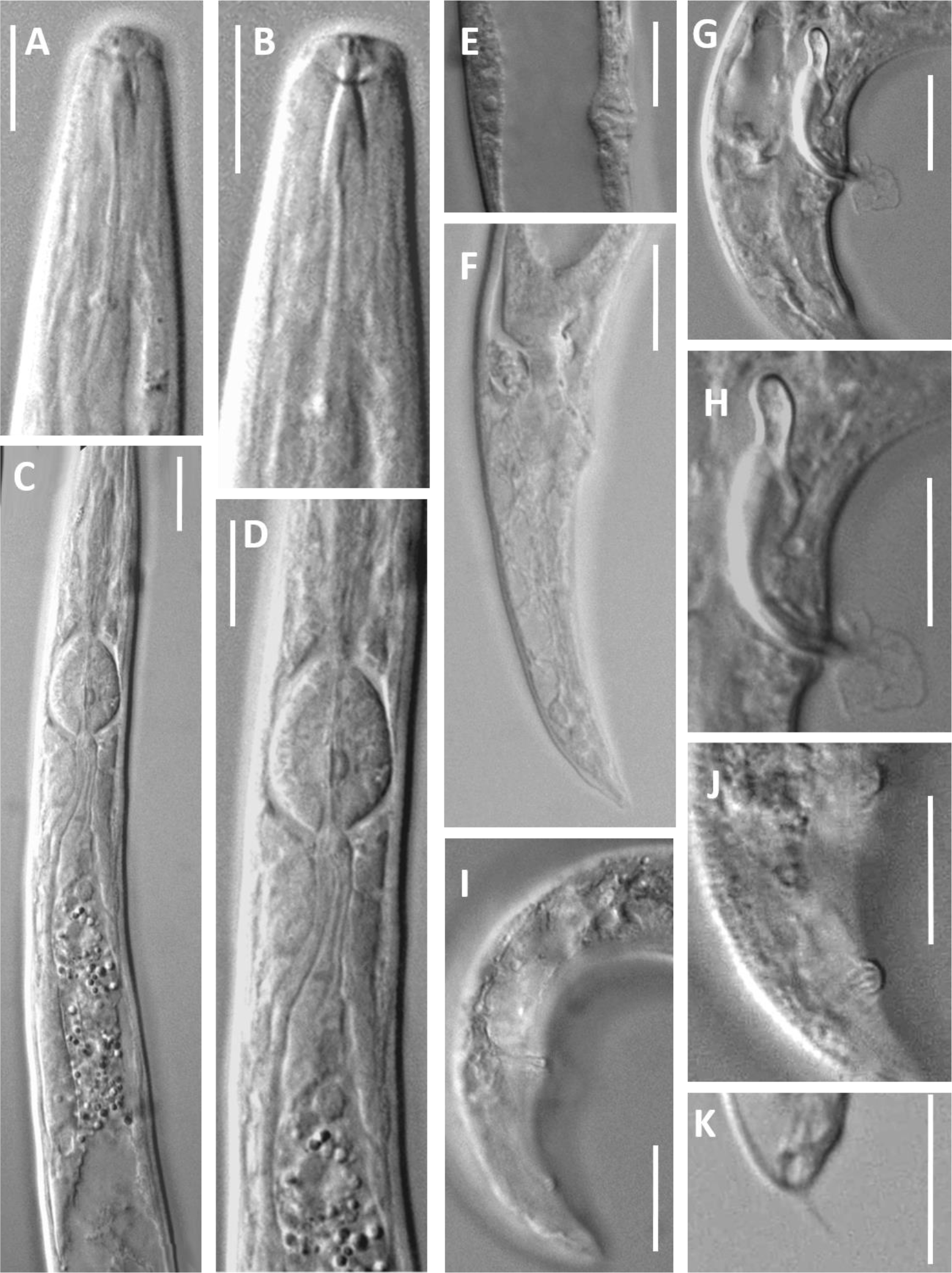
Photomicrographs of *Ficophagus glomerata* n. sp., anterior end of a male showing a stylet and basal knob (A – B) and a distinct median bulb and pharyngeal glands (C – D). Vulval region of female showing vulval opening (E), and tail region of the male showing anal opening (F), spicule (G – H), genital papillae (I – J) and spicate tip (K). Scale bar: (A – K) 10 µm.

**Fig. 3.**
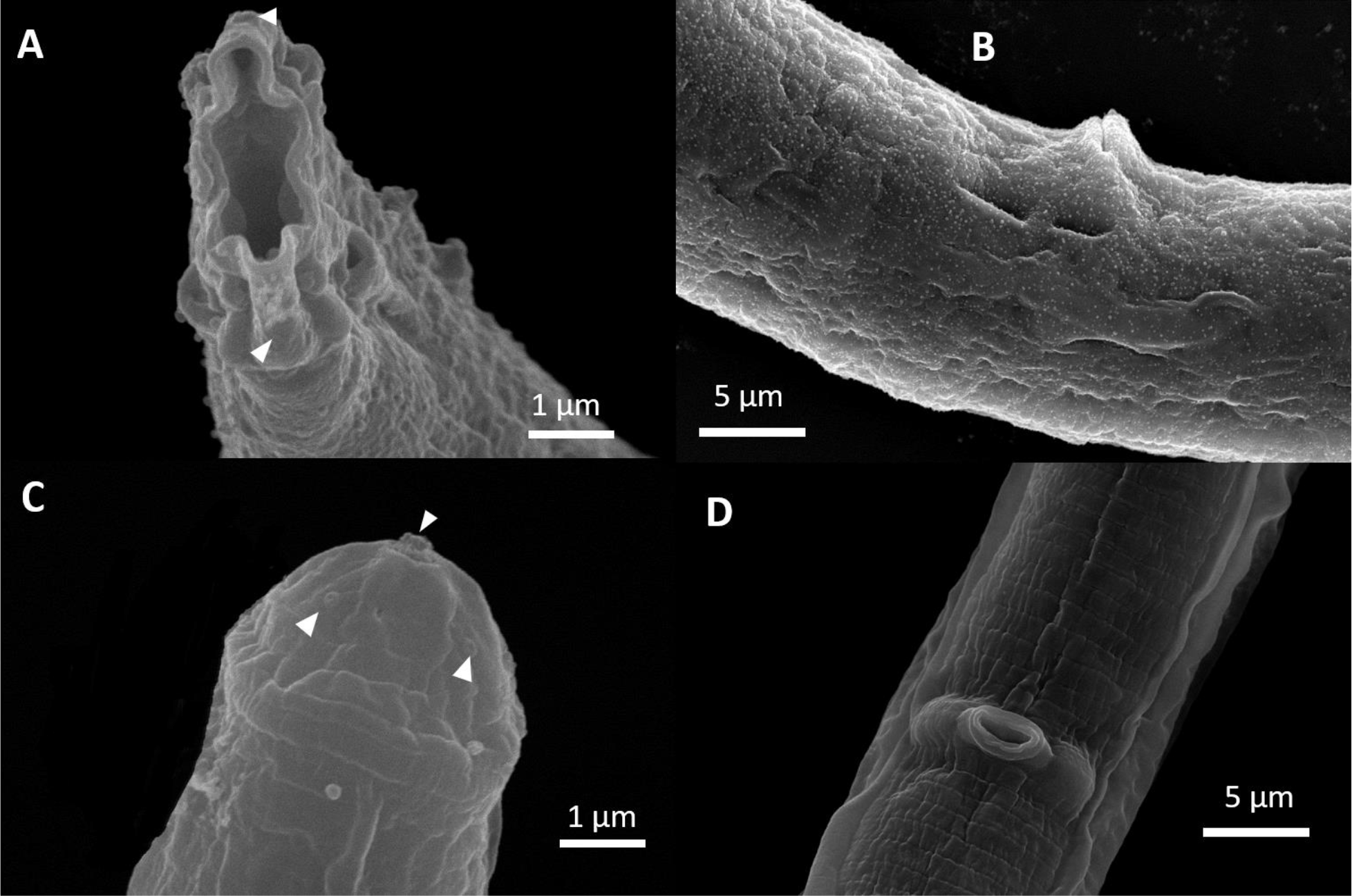
SEM observations: A: Anterior end; B: Vulval region of female of *Teratodiplogaster glomerata* n. sp.; C: Anterior end; D: Vulval region of female *Ficophagus glomerata* n. sp. (arrows indicate points to the amphidal pore and cephalic sensilla).

**Fig. 4.**
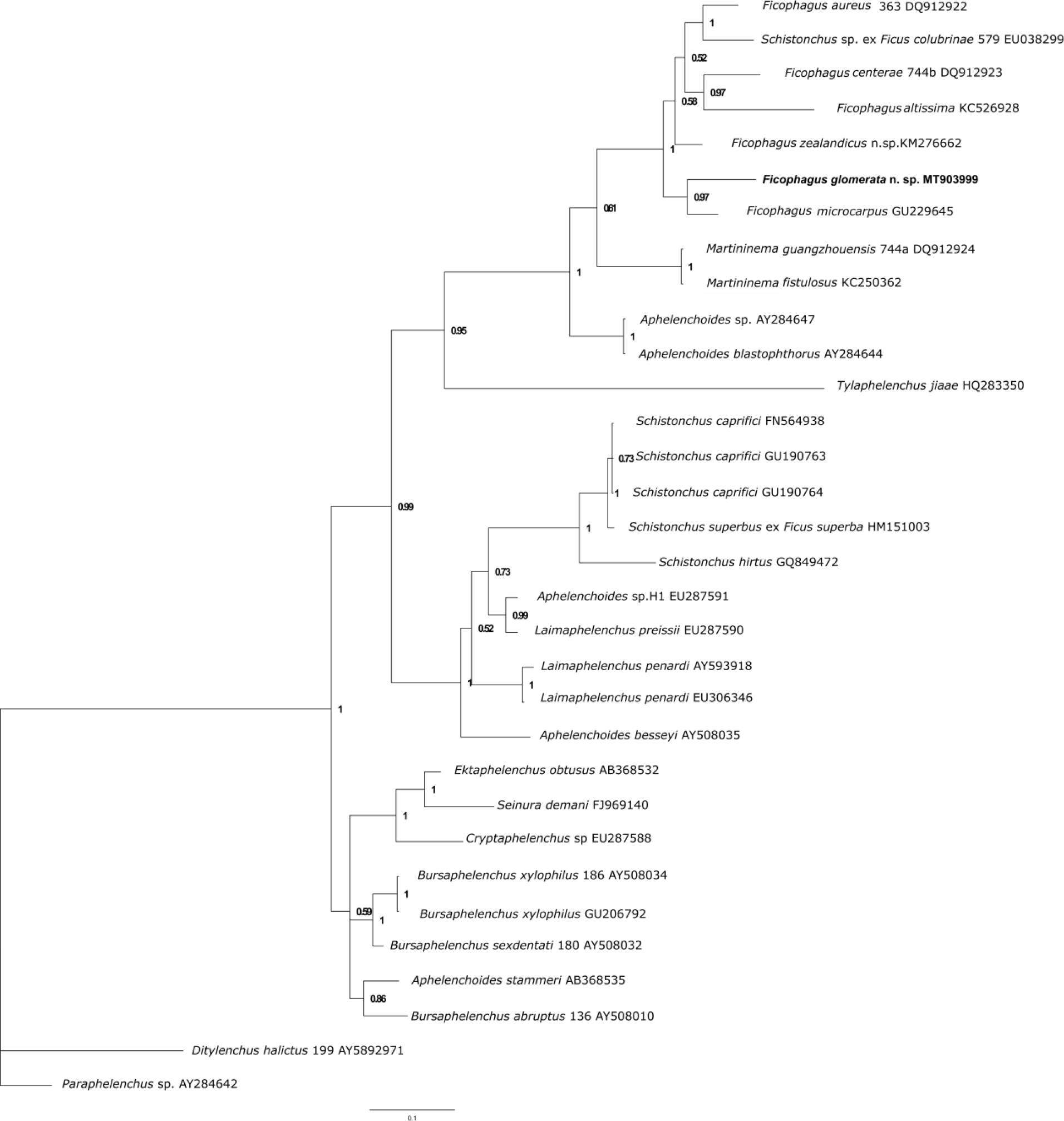
The Bayesian tree inferred from the 18S gene for *Ficophagus glomerata* n. sp. under the GTR + I + G model (lnL = 5957.9728; freqA = 0.2547; freqC = 0.1966; freqG = 0.2654; freqT = 0.2833; R (a) = 0.0959; R (b) = 0.2182; R (c) = 0.1305; R (d) = 0.0852; R (e) = 0.4025; R (f) = 0.0676; Pinva = 0.27; Shape = 0.593). The accession numbers of the compared sequences are indicated in the form: SSU. Posterior probability values exceeding 50% are given on appropriate clades.

**Fig. 5.**
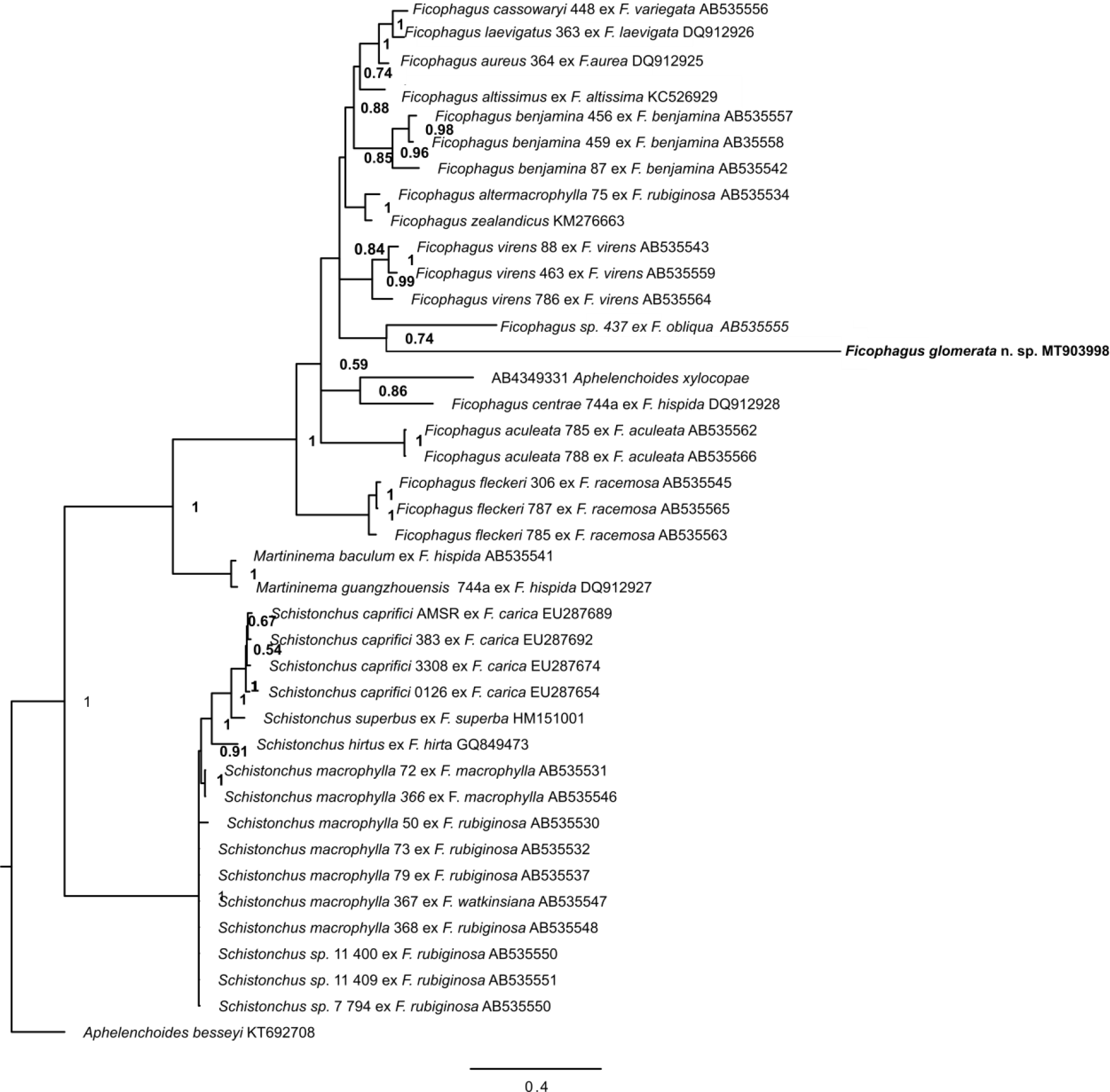
The Bayesian tree inferred from the 28S gene for *Ficophagus glomerata* n. sp. under the GTR + I + G model (lnL = 9439.3496; freqA = 0.2437; freqC = 0.1807; freqG = 0.2957; freqT = 0.28; R (a) = 0.0592; R (b) = 0.2708; R (c) = 0.1709; R (d) = 0.0879; R (e) = 0.397; R (f) = 0.0772; Pinva = 0.0117; Shape = 0.7021). The accession numbers of the compared sequences are indicated in the form: D2-D3 LSU. Posterior probability values exceeding 50% are given on appropriate clades.

MEASUREMENTS. Table 1, 2.

**Table 1.** Morphometric characteristics of *Ficophagus glomerata* n. sp. All measurements are in μm and in the form: mean ± s.d. (range).

| Character | *Ficophagus glomerata* n. sp. |  |  |
| --- | --- | --- | --- |
|  | Male |  | Female |
|  | Holotype | Paratypes | Paratypes |
| n |  | 9 | 10 |
| L | 525 | 567 ± 58<br>(469-656) | 761 ± 81<br>(604-886) |
| a | 25 | 26.5 ± 2.4<br>(23.4-30.9) | 24.0 ± 2.3<br>(20-28) |
| b | 7.3 | 7.4 ± 0.3<br>(6.7-7.8) | 7.6 ± 1.4<br>(5.2-9.6) |
| c | 17.2 | 16.6 ± 2.6<br>(12-20) | 20.7 ± 1.6<br>(18-23) |
| c' | 1.8 | 2.0 ± 0.4<br>(1.5-2.7) | 1.6 ± 0.2<br>(1.3- 1.9) |
| T or V (%) | 76.9 | 72 ± 3.3<br>(65-77) | 68.6 ± 4.9<br>(61.2-76.7) |
| Max. body diam. | 20.9 | 21.3 ± 1.0<br>(19-23) | 31.4 ± 0.7<br>(30.2-32.1) |
| Stylet length | 21.2 | 21.3 ± 0.6<br>(20-22) | 22.0 ± 0.9<br>(20-23) |
| Median bulb (length) | 16.6 | 16.2 ± 0.6<br>(14-17) | 14.3 ± 0.9<br>(12.4 -15.6) |
| Median bulb (width) | 10.6 | 10.6 ± 0.3<br>(10-11) | 10.4 ± 0.5<br>(10-11) |
| Ant.enf to SE pore | 14 | 14 ± 0.7<br>(13-15) | 14.5 ± 0.8<br>(13-16) |
| Ant. end to base of metacarpus | 55 | 59 ± 5.5<br>(51-68) | 64.6 ± 2.3<br>(62-69) |
| Ant. end to base of pharyngeal gland | 114 | 80.4 ± 12<br>(68-114) | 101.5 ± 15<br>(78-114) |
| Nerve ring from anterior end | 68 | 70.8 ± 6<br>(62-82) | 77.9 ± 6.1<br>(71-87.4) |
| Post uterine sac | - | - | 6.0 ± 0.3<br>(5.4-6.3) |
| Tail length | 35.1 | 33.5 ± 6.6<br>(23.6-42) | 38.9 ± 2.1<br>(34.5-42.3) |
| Spicule length (chord) | 16.6 | 16.8 ± 1<br>(14-17) | |

**Table 2.** Comparative morphometric data of *Ficophagus glomerata* n. sp., other congeners and two *Schistonchus* sp. described from India, associated with *Ficus racemosa*. All measurements are in µm and in the form mean ± s.d. (range).

| Character | *S. racemosa* |  | *S. osmani* |  | *F. aculeate* |  | *F. fleckeri* |  | *F. altermacrophylla* |  | *F. cuculloracemosus* |  | *F. flagelloracemosus* |  | *F. glomerata* n. sp. * |  |
| --- | --- | --- | --- | --- | --- | --- | --- | --- | --- | --- | --- | --- | --- | --- | --- | --- |
|  | Male | Female | Male | Female | Male | Female | Male | Female | Male | Female | Male | Female | Male | Female | Male | Female |
|  | Paratypes | Paratypes | Paratypes | Paratypes | Paratypes | Paratypes | Paratypes | Paratypes | Paratypes | Paratypes | Paratypes | Paratypes | Paratypes | Paratypes | Paratypes | Paratypes |
| n | 35 | 35 | 30 | 30 | 15 | 14 | 22 | 27 | 12 | 10 | 11 | 14 | 12 | 15 | 9 | 10 |
| L | 653<br>(500-660) | 792<br>(600-800) | (700-840) | (718-910) | 450 ± 33<br>(396-523) | 619 ± 35<br>(548-671) | 547 ± 32<br>(498-609) | 727 ± 78<br>(572-889) | 560 ± 44<br>(445-618) | 500 ± 49<br>(411-571) | 523.6±56.1<br>(441-609) | 785.7±57.5<br>(707-906) | 379.8±28.6<br>(342-446) | 475.9±18.3<br>(442-504) | 567 ± 58<br>(469-656) | 761 ± 81<br>(604-886) |
| a | 26<br>(24-26) | 117.2<br>(14-21) | (26-30) | (27-34) | 27 ± 2.8<br>(23-33) | 18 ± 2.2<br>(14-23) | 30 ± 1.5<br>(27-33) | 23 ± 3.9<br>(15-35) | 28 ± 2<br>(25-31) | 22 ± 3<br>(17-29) | 26.6±1.57<br>(24.5-29.3) | 22.1±2.81<br>(17.6-27.8) | 27.3±3.3<br>(22.8-32.9) | 26.9±1.52<br>(24.3-30.4) | 26.5 ± 2.4<br>(23.4-30.9) | 24.0 ± 2.3<br>(20-28) |
| b | 5.4<br>(5.2-6.2) | (5.2-6.2) | (10-13) | (11.5-13.5) | 3.3 ± 0.2<br>(2.6-4.0) | 4.9 ± 0.2<br>(4.7-5.1) | 5.5 ± 0.5<br>(4.8-6.4) | 7.2 ± 0.3<br>(7.0-7.4) | 4 ± 0.6<br>(3-5) | 5 ± 1.2<br>(3-8) | 7.9±0.88<br>(6.0-9.5) | 12.3±1.12<br>(10.9-15.1) | 9.1±0.84<br>(8.1-11.2) | 9±0.54<br>(8.0-10.1) | 7.4 ± 0.3<br>(6.7-7.8) | 7.6 ± 1.4<br>(5.2-9.6) |
| c | 28<br>(22-28) | 21.6<br>(18-22) | (12-15) | (14.8-19.6) | 12.2 ± 1.0<br>(11-14) | 19.3 ± 2.0<br>(17-23) | 18 ± 1.5<br>(16-21) | 21 ± 3.4<br>(12-29) | 15 ± 1<br>(13-16) | 17 ± 2<br>(14-21) | 17.6±1.3<br>(16-20.5) | 16.9±2.36<br>(14.5-21.1) | 24.3±2.1<br>(21.1-27.5) | 13.9±1.1<br>(12.8-16.8) | 16.6 ± 2.6<br>(12-20) | 20.7 ± 1.6<br>(18-23) |
| c' | - | - | - | - | 2.3 ± 0.2<br>(1.8-2.8) | 2.4 ± 0.4<br>(2.0-3.1) | 1.9 ± 0.2<br>(1.6-2.4) | 2.6 ± 0.7<br>(1.6-4.6) | 2 ± 0.2<br>(1.7-2.4) | 3 ± 0.3<br>(2.8-3.8) | 1.8±0.14<br>(1.7-1.9) | 3.4±0.5<br>(2.7-4.1) | 1.3±0.09<br>(1.2-1.4) | 4.5±0.48<br>(3.4-5.2) | 2.0 ± 0.4<br>(1.5-2.7) | 1.6 ± 0.2<br>(1.3- 1.9) |
| T or V (%) | 31<br>(30-48) | 60<br>(60-76) | - | (78-84) | 55 ± 1<br>(48-57) | 75 ± 1.2<br>(73-77) | 41 ± 13.4<br>(22-80) | 76 ± 4.3<br>(67-86) | 330 ± 49<br>(273-401) | 85 ± 9<br>(70-95) | - | 73.2±2.0<br>(70.1-77.6) | - | 76.2±0.96<br>(74.7±78.2) | 72 ± 3.3<br>(65-77) | 68.6 ± 4.9<br>(61.2-76.7) |
| Max. body diam. | - | - | (25-28.5) | (21-27) | 17 ± 1<br>(15-19) | 34 ± 3<br>(29-40) | 18 ± 1.5<br>(15-19) | 33 ± 7<br>(19-54) | 18 ± 2<br>(15-19) | 23 ± 2<br>(19-25) | - | - | - | - | 21.3 ± 1.0<br>(19-23) | 31.4 ± 0.7<br>(30.2-32.1) |
| Stylet length | - | 23<br>(21-24) | (19-21) | (21-27) | 15 ± 2<br>(13-18) | 14 ±1.5<br>(13-17) | 17 ± 0.8<br>(15-17) | 18 ± 1.5<br>(15-19) | 22 ± 3<br>(16-25) | 21 ± 2<br>(17-26) | 22<br>(22) | 22<br>(22) | 14<br>(14) | 14<br>(14) | 21.3 ± 0.6<br>(20-22) | 22.0±0.9<br>(20-23) |
| Median bulb (length) | 17<br>(13-17) | 17<br>(13-17) | (10-12) | (12-13) | - | - | - | - | - | - | - | - | - | - | 16.2 ± 0.6<br>(14-17) | 14.3 ± 0.9<br>(12.4 -15.6) |
| Median bulb (width) | 13<br>(10-13) | 15<br>(12-15) | (8-9) | (8-9) | - | - | - | - | - | - | - | - | - | - | 10.6 ± 0.3<br>(10-11) | 10.4 ± 0.5<br>(10-11) |
| Ant. end to Ex pore | - | - | - | - | 6 ± 2.0<br>(4-9) | 6 ± 1.1<br>(5-8) | 10 ± 1.4<br>(8-11) | 8.0 ± 0.6<br>(7-8) | 6 ± 1<br>(3-7) | 6 ± 2<br>(5-9) | 10±1.89<br>(7-12) | 10.1±1.8<br>(7-14) | 16.7±2.0<br>(12-19) | 11.4±0.44<br>(11-13) | 14 ± 0.7<br>(13-15) | 14.5 ± 0.8<br>(13-16) |
| Ant. End<br>to base<br>metacarpus | - | - | - | - | 53 ± 3<br>(49-58) | 54 ± 7<br>(47-77) | 66 ± 3.7<br>(60-73) | 58 ± 9.5<br>(41-87) | 65 ± 7<br>(60-71) | 69 ± 6<br>(55-77) | - | - | - | - | 59 ± 5.5<br>(51-68) | 64.6 ± 2.3<br>(62-69) |
| Ant. End<br>to base<br>pharyngeal<br>gland | - | - | - | - | 135 ± 7<br>(123-154) | 127 ± 5<br>(117-135) | 100 ± 6<br>(87-110) | 119 ± 11<br>(112-127) | 130 ± 20<br>(112-163) | 10 ± 20<br>(114-168) | - | - | - | - | 80.4 ± 12<br>(68-114) | 101.5 ± 15<br>(78-114) |
| Nerve ring<br>from<br>anterior<br>end | - | - | - | - | 60 ± 3<br>(53-65) | 57 ± 4<br>(52-63) | 72 ± 4.5<br>(65-78) | 66 ± 3<br>(61-70) | 72 ± 8<br>(60-85) | 76 ± 8<br>(60-83) | - | - | - | - | 70.8 ± 6<br>(62-82) | 77.9 ± 6.1<br>(71-87.4) |
| Post<br>uterine sac | - | - | - | - | - | 6 ± 1.5<br>(4-9) | - | 18 ± 8<br>(6-39) | - | 14 ± 3<br>(8-20) | - | - | - | - | - | 6.0 ± 0.3<br>(5.4-6.3) |
| Tail length | - | - | - | - | 37 ± 4<br>(31-43) | 32 ± 3<br>(28-36) | 30 ± 3<br>(26-38) | 37 ± 8<br>(24-62) | 34 ± 2<br>(28-37) | 29 ± 4<br>(23 -34) | 29.7±2.1<br>(25-32) | 47.3±7.58<br>(35-62) | 15.7±0.86<br>(14-16) | 34.3±2.58<br>(30-38) | 33.5 ± 6.6<br>(23.6-42) | 38.9 ± 2.1<br>(34.5-42.3) |
| Spicule<br>length<br>(chord) | - | - | (37-39.5) | - | 16 ± 1<br>(15-17) | - | 17 ± 0.9<br>(16-18) | - | 21 ± 2<br>(18-24) | - | 21.4±0.67<br>(20-22) | - | 10.1±0.5<br>(9-11) | - | 16.8 ± 1<br>(14-17) | - |

DESCRIPTION

### Male

Body slender, small- to medium-sized, C-shaped when heat-killed, tapering at both ends with maximum width at 2/3^rd^ of the body length from anterior end. Cuticle with annules < 1 µm wide at mid body. Lateral fields with single thick continuous ridge representing two lines. Lip region off-set; cephalic framework moderately sclerotized. Oral aperture surrounded by a sclerotized circular rim; labial disc absent; labial and cephalic sensilla papilliform, not discernible in LM but visible in SEM. Stylet 20–22 µm long, robust with small, rounded / sloping knobs; conus 60-65% of stylet length. Procorpus 35.0-51.7 µm long, median bulb round to ovoid, muscular, separated from isthmus by a constriction; isthmus short, indistinguishable from elongated basal overlap represented by a long dorsal and small ventro-lateral lobe. Secretory-excretory pore located in anterior region, 13-16 *μ*m or *ca* 1.2-1.5 lip region diam. from anterior end. Pharyngo-intestinal junction posterior to median bulb in isthmus region. Deirids, hemizonid and phasmids not discernible. Testis usually on left side of intestine; seminal vesicle having amoeboid sperms. Spicules slender, strongly arcuate, with hammer-shaped capitulum with a depression comprising an ovoid, spatulate condylus and a long digitate rostrum. Gubernaculum absent. Three pairs of genital papillae in configuration of P1 subventral adcloacal pair and P2 subventral, halfway between cloaca and tail tip and P3 about 3-4 μm anterior to tail terminus. Phasmids located laterally between P1 and P2. Tail conoid, strongly ventrally curved ending into a fine bristle-like spike. Bursa absent.

### Female

Body slender, ventrally curved or C-shaped when heat-killed, tapering at both ends with maximum diam. at mid-body. Lateral fields with a single ridge as observed under SEM occupying ∼2 µm width; labial sensilla papilliform surrounding oral opening; four cephalic papillae present. Cephalic framework moderately sclerotized, Amphids not discernible under LM. Stylet moderately built with well-developed, rounded/ sloping knobs; conus approximately 60-65% of stylet length. Procorpus slender (46.4 µm-55.6 µm) long; median bulb well-developed, ovoid in shape, with valve plates slightly posterior to middle. Pharyngeal glands and pharyngo-intestinal junction similar to those observed in male. Deirids and hemizonid not seen. Excretory pore located in anterior region, *ca* 1.2-1.5 lip region diam. from anterior end. Reproductive system monodelphic-prodelphic. Ovary large, outstretched or with flexure; reflexed part containing oocytes usually in single tier in the proximal region. Oviduct narrow, spermatheca not differentiated; uterus slightly spacious chamber; post-uterine sac 0.4-0.8 times vulval body diam.; Vulva post-equatorial; vulval lips protruding in some specimens. Anus a crescent-shaped slit. Tail ventrally arcuate, four-times anal body diam. long with an obtuse or mucronate tip.

#### TYPE HABITAT AND LOCALITY

*Ficophagus glomerata* n. sp. was collected from syconia of *Ficus racemosa* in and around the campus of Indian Institute of Science, Bangalore, Karnataka, India at coordinates 13.0219° N, 77.5671° E.

#### TYPE SPECIMENS

One holotype male (*F. glomerata* n. sp./1), eight paratype males (*F. glomerata* n. sp./2-8) and nine paratype females (*F. glomerata* n. sp./1-9) of *F. glomerata* n. sp. on slides were deposited in Indian Institute of Science, Bangalore, Karnataka, India. One female paratype (*F. glomerata* n. sp./10) and one male paratype (*F. glomerata* n. sp./9) were deposited in the National Nematode Collection, Indian Agricultural Research Institute, New Delhi.

**Etymology:** The species name “*glomerata*” derived from *Ficus glomerata,* is a synonym of *Ficus racemosa*.

#### DIAGNOSIS AND RELATIONSHIP

The new species *F. glomerata* n. sp. can be characterised by small body having ‘b’=5.2-9.6, ‘c’= 18-23; slightly setoff lip region having well developed cephalic framework; slender stylet with small, rounded/ sloping knobs; anteriorly located excretory pore; ovoid median bulb with relatively posteriorly-placed valve plates; males with sickle-shaped spicules having hammer-shaped capitulum, represented by an ovoid spatula-shaped condylus and long digitate rostrum and tail conoid with fine, hair-like terminal spike and three pairs of genital papillae.

*Ficophagus glomerata* n. sp. closely resembles *F. fleckeri* (Davies *et al*., 2013) Davies & Bartholomaeus, 2015 in most morphometric and morphological characteristics but differs in having less lines (2 *vs* 4) in lateral fields; greater ‘b’ value in male (6.7-7.8 *vs* 4.8-6.4); larger stylet (20-23 μm *vs* 15-19 μm); spicules with an ovoid, spatula-shaped condylus, and rostrum digitate and separate (*vs* prominent condylus with rostrum merging into ventral arm); male tail with 3-4 μm hair-like spike (*vs* 2-3 μm long conical terminal mucro) and female tail with a narrow blunt mucro or terminus (*vs* acutely pointed terminus in *F. fleckeri*).

The new species closely resembles *F. microcarpus* Zeng *et al*., 2011, in most morphometric and morphological characteristics but differs in relatively smaller ‘b’ (5.2-9.6 *vs* 8.5-13.0), ‘ć’ (1.3-1.9 *vs* 2.6-3.6) and greater ‘c’ (18-23 *vs* 12.3-16.6) values in females; relatively posteriorly located excretory pore (13-16 μm *vs* 3.5-5.5 μm) from anterior end; amphids indiscernible (*vs* prominent); post-uterine sac longer (more than vulval body diam. *vs* 0.4-0.8 vulval body diam.); males are relatively larger (604-886 μm *vs* 400-486 μm) with sickle-shaped spicules having an ovoid, spatulate (*vs* small, rounded) condylus and long, digitate (*vs* narrowly rounded to short digitate) rostrum and cucullus absent (*vs* present at distal end of spicules in *F. microcarpus*).

*Ficophagus glomerata* n. sp. differs from *F. virens* Bartholomaeus *et al*., 2009 in having lateral fields with 2 (*vs* 4) lateral lines; females with relatively smaller ‘a’ (20-28 *vs* 28-34) and ‘c’ (18-23 *vs* 19-41) values, longer stylet (20-23 µm *vs* 14-16 µm); post-uterine sac longer (more than corresponding vulva body diam. *vs* 0.6-0.7 vulval body diam.); males having greater ‘b’ (6.7-7.8 *vs* 3-5) value, sickle-shaped spicules having long, digitate (*vs* short, rounded to digitate) rostrum and tail with hair-like terminal spike (*vs* without terminal mucro in *F. virens*).

The new species differs from *F. altermacrophylla* Lloyd & Davies, 1997 in having lateral fields with 2 (*vs* 3) lateral lines; larger females (604-886 µm *vs* 411-571 µm) with smaller ‘ć’ (1.3-1.9 *vs* 2.8-3.8), relatively posteriorly located (13-16 µm *vs* 5-9 µm) excretory pore, longer stylet (20-23 µm *vs* 14-18 µm); and relatively anteriorly located vulva (V= 61.2-76.7 *vs* 70-95); males having greater body diam. (19-23 µm *vs* 15-19 µm) and ‘b’ value (5.2-9.6 *vs* 3-5); posteriorly located (13-16 µm *vs* 3-7 µm) excretory pore; post-uterine sac longer (more than corresponding vulval body diam. *vs* 0.4-0.8 vulval body diam.); and smaller (14-17 µm *vs* 18-24 µm), sickle-shaped spicules with an ovoid, spatulate condylus and long, digitate rostrum (*vs* rose-thorn-shaped spicules with knob-like condylus and short, angular rostrum in *S. altermacrophylla*). *Ficophagus glomerata* n. sp. was compared with all the species of *Ficophagus* reported from India. The earlier reported species, *S. racemosa*, *S. osmani* isolated from *Ficus racemosa* were considered *species inquirendae* (Davies *et al*., 2015) along with *S. hispida* (Kumari & Reddy, 1984) isolated from *Ficus hispida,* due to insufficient information particularly on the position of the excretory pore, spicules and position of the caudal papillae. The present species, however, can be well differentiated from *S. racemosa, S. osmani* and *S. hispida* on account of morphometric values including shorter (*vs* larger) post-uterine sac in females; amoeboid (*vs* flagellate/ rod-shaped) sperms; smaller (*vs* longer) spicules and three pairs (*vs* two pairs) of genital papillae.

Bajaj and Tomar (2014) published the description of eight species of *Schistonchus* isolated from three species of *Ficus*. Six of them appear to belong to genus *Ficophagus* largely on the position of excretory pore although the differentiating characters of the species are not clear. Only two species *S. flagellobenghalensus* (Bajaj & Tomar, 2014)*, S. mucroracemosus* with a posterior excretory pore do not fit into *Ficophagus*. In absence of molecular characterization, the status of the species could not be confirmed due to considerable morphological fluidity and morphometric overlap. Nevertheless, to verify the status and distinctness of *F. glomerata* n. sp., comparisons were made with species described from India.

The new species differs from *F.* (= *Schistonchus*) *mucrobenghalensus* (Bajaj & Tomar, 2014) n. comb. in having relatively smaller ‘b’ (6.7-7.8 *vs* 7.2-9.1) and ‘c’ (12-20 *vs* 21.9-26.7) values; dissimilar larger spicules; gubernaculum absent (*vs* faintly visible); genital papillae three pairs (*vs* four pairs) and tail with hair-like spike/mucro (*vs* tail devoid of mucro) in males and smaller ‘b’ value (5.2-9.6 *vs* 7.0-9.2) in females and tail with short, blunt (*vs* long, pointed terminal) mucro in both sexes and genital papillae in males three pairs [*vs* four pairs in *F. mucrobenghalensus*]. A disparity in values of stylet length and excretory pore position was observed between description (page no. 199) and table (3).

The new species differs from *F.* (= *Schistonchus*) *antherobenghalensus* (Bajaj & Tomar, 2014) n. comb. in having longer stylet (20-22 µm *vs* 15 µm); dissimilar spicules; and tail with hair-like spike (*vs* tail devoid of mucro) in male and tail in females having short, blunt (*vs* long, pointed terminal) mucro in *F. antherobenghalensus*. A disparity in position of excretory pore as stated in description (page no. 200) and that given in table (4) was found.

*Ficophagus glomerata* n. sp. differs from *F.* (= *Schistonchus*) *flagelloracemosus* in having smaller ‘b’ (6.7-7.8 *vs* 8.1-11.2), ‘c’ (12-20 *vs* 21.1-27.5) and greater ‘ć’ (1.5-2.7 *vs* 1.2-1.4) values in males and smaller ‘ć’ value (1.3-1.9 *vs* 3.4-5.2) in females; longer stylet (20-22 µm *vs* 14 µm); males with larger (14-17 µm *vs* 9-11 µm) spicules and gubernaculum absent [*vs* faintly visible in *F. flagelloracemosus* n. comb.]

The new species differs from *F.* (= *Schistonchus*) *cuculloracemosus* n. comb.in having smaller ‘b’ (5.2-9.6 *vs* 10.9-15.1) value in females; stoma with rounded (*vs* elongated) knobs; excretory pore relatively posteriorly located (13-15 µm *vs* 7-12 µm) in males; spicules smaller (14-17 µm *vs* 20-22 µm), without (*vs* with cucullus) at distal end in *F. cuculloracemosus* n. comb.

The morphometric differences with the closely related species of *Ficophagus* reported from *Ficus racemosa* have been given in the Table 2.

### Molecular phylogenetic relationship

Partial SSU and D2-D3 of LSU genes were sequenced for *F. glomerata* n. sp. The relative placement of *F. glomerata* n. sp. among the other sequenced *Ficophagus* and *Schistonchus* species was analyzed. The Bayesian tree (Fig. 4) for SSU was constructed using *Ditylenchus halictus* (Zhao *et al*., 2015) as an outgroup and suggested that: i) *F. glomerata* n. sp. forms a sister species with *S. microcarpus* having 97% posterior probability, ii) the genera *Aphelenchoides* Fischer, 1894, *Bursaphelenchus* Fuchs, 1937, *Laimaphelenchus* Fuchs, 1937 or *Schistonchus* are not monophyletic which stands in concordance with earlier studies (Zhao *et al*., 2015). The Bayesian tree inferred from D2-D3 of LSU genes (Fig. 5) using *Aphelenchoides besseyi* (Zhao *et al*., 2015) as an outgroup, to analyze the relationships of the species in the genus *Schistonchus* suggested that: i) all the sequenced *Schistonchus* species are divided into two clades with 100% posterior probability support in accordance with earlier work done on the phylogeny of S*chistonchus* (*= Ficophagus*) species (Zeng *et al*., 2011, Zhao *et al*., 2015); ii) *F. glomerata* n. sp. appeared to be closest to an isolate of *Ficophagus* from *Ficus obliqua* in Australia, and forms a clade with 74% posterior probability support; this forms a sister group to two other clades, one formed by *F. virens* and another one which includes *F. zealandicus*, *F. altermacrophylla* and *F. benjamina* with 84% posterior probability support. Accession numbers in form of SSU/D2-D3 LSU for *Ficophagus glomerata* n. sp. are MT903999/ MT903998.

***Teratodiplogaster glomerata* n. sp.**

(Figs. 3, 6-8)

**Fig. 6.**
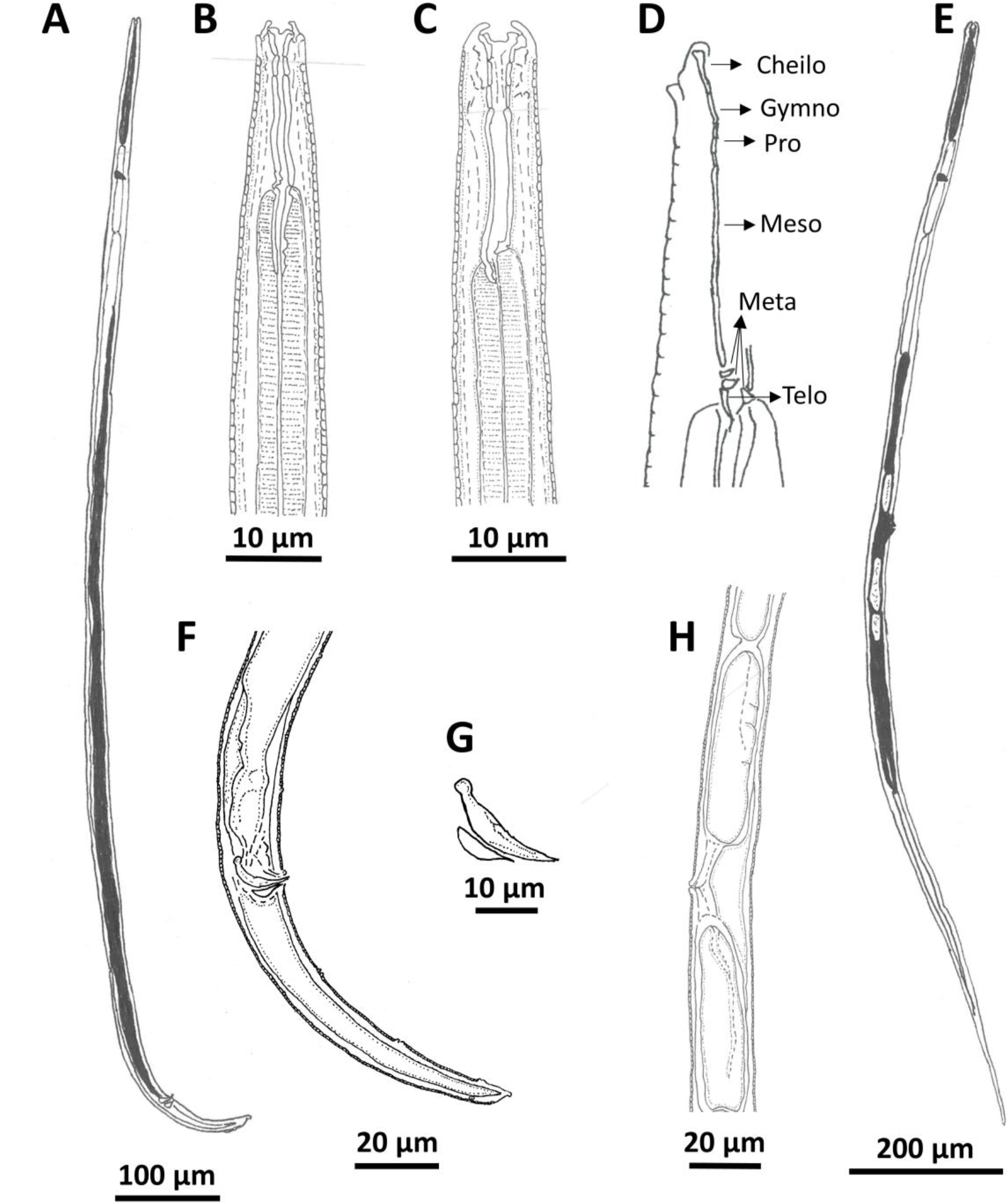
*Teratodiplogatser glomerata* n. sp. lateral view (A) Habitus of male, (B) Anterior region of male, (C) Anterior region of female, (D) Schematic representation of lateral view of stomatal morphology, (E) Habitus of male, (F) Tail region of male, (G) Spicule and gubernaculum and (H) Vulval region of female.

**Fig. 7.**
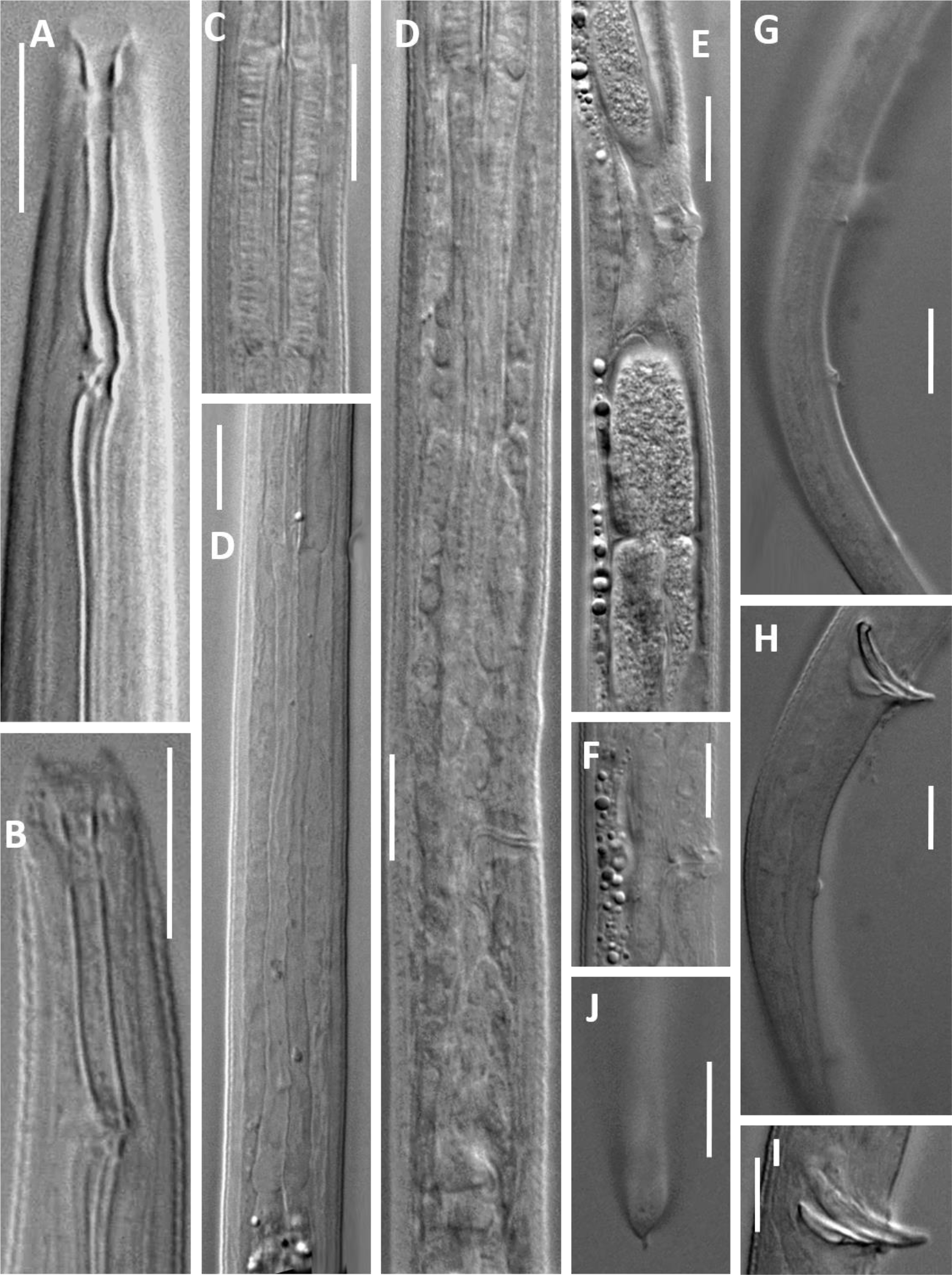
Photomicrographs of *Teratodiplogaster glomerata* n. sp. (A, B) Head morphology of a male and female respectively. Anterior and posterior pharynx (C – D), (E) Female gonadal region showing vulva region showing the vulval opening and the eggs, (F) vulval opening, (G – H) Tail region of a male at different focal planes showing genital papillae, (I) the male spicule, and (J) the mucronated tip. Scale bar: (A – J) 10 µm.

**Fig. 8.**
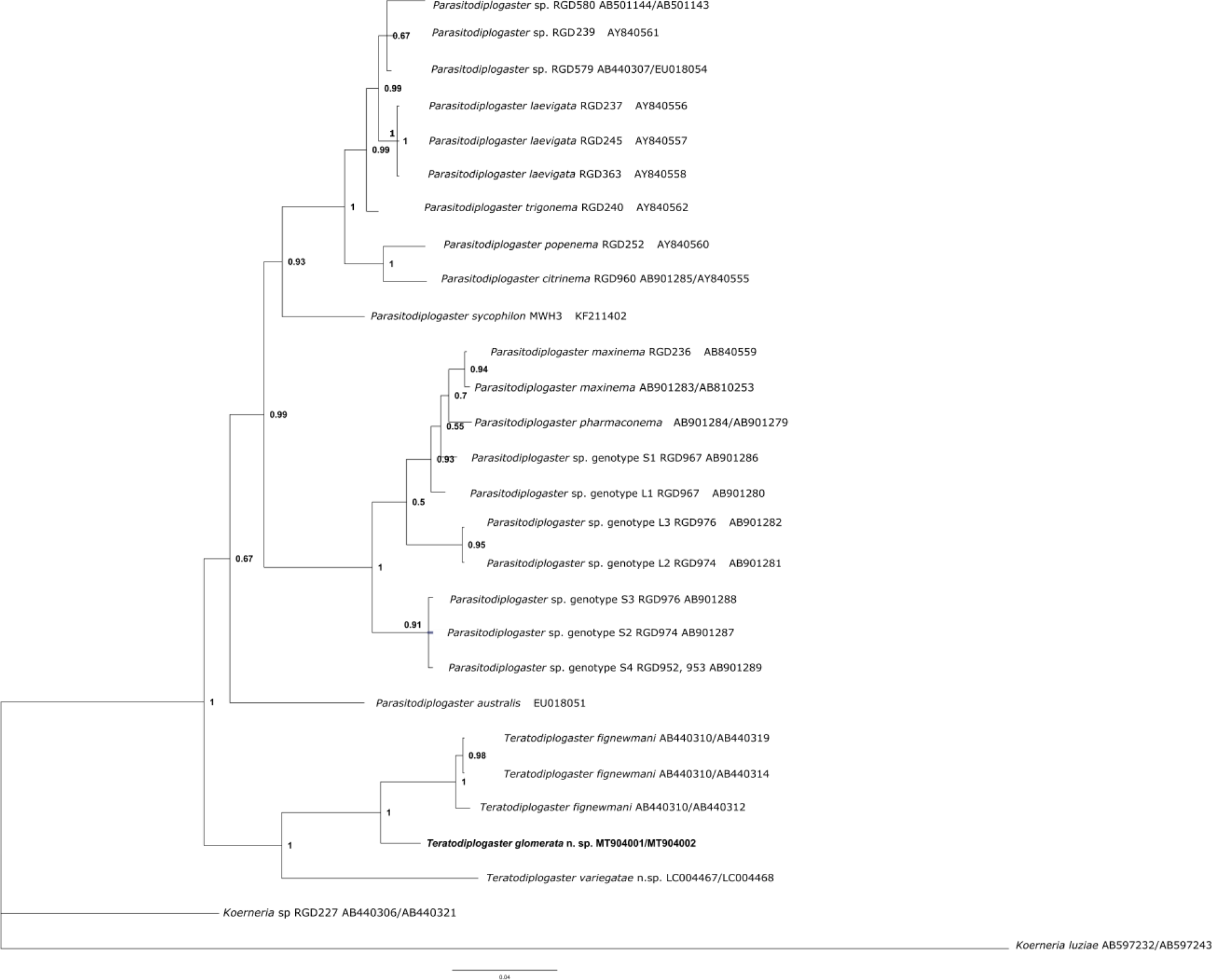
The Bayesian tree inferred from the 18S and 28S gene for *Teratodiplogaster glomerata* n. sp. under the GTR + I + G model (lnL = 6263.9613; freqA = 0.1647; freqC = 0.2377; freqG = 0.336; freqT = 0.2617; R (a) = 0.0565; R (b) = 0.2121; R (c) = 0.1209; R (d) = 0.0504; R (e) = 0.4818; R (f) = 0.0783; Pinva = 0.251; Shape = 0.915). The accession numbers of the compared sequences are indicated in the form: SSU/D2-D3 LSU. Posterior probability values exceeding 50% are given on appropriate clades.

MEASUREMENTS. Table 3

**Table 3.** Comparative morphometric data of *Teratodiplogaster glomerata* n. sp. and other congeners associated with *Ficus racemosa*. All measurements are in μm and in the form: mean ± s.d. (range).

| Character | *T. variegatae* |  | *T. fignewmani* |  | *T. martini* |  | *T. racemosus* |  | *T. glomerata* n. sp. |  |  |
| --- | --- | --- | --- | --- | --- | --- | --- | --- | --- | --- | --- |
|  | Male | Female | Male | Female | Male | Female | Male | Female | Male |  | Female |
|  | Paratypes | Paratypes | Paratypes | Paratypes | Paratypes | Paratypes | Paratypes | Paratypes | Holotype | Paratypes | Paratypes |
| n | 9 | 10 | 10 | 11 | 10 | 13 | 9 | 13 |  | 9 | 10 |
| L | 1019 ± 117<br>(854-1283) | 965 ± 120<br>(805-1173) | 2119 ± 311<br>(1850-2700) | 2705 ± 636<br>(2160-3895) | 2171 ± 267<br>(1825-2650) | 2253 ± 325<br>(1750-3100) | 1771.1 ± 295.6<br>(1412-2184) | 2030.9 ± 236.2<br>(1587-2444) | 1329 | 1271 ± 48<br>(1195-1345) | 1506 ± 83<br>(1388-1657) |
| a | 40.9 ± 4.6<br>(29.9-47.4) | 30.7 ± 5.8<br>(26.1-45.6) | 67.8 ± 6.6<br>(52.9-75.9) | 79.9 ± 10.6<br>(59.0-92.0) | 62.6 ± 4.6<br>(52.6-70.0) | 67.1 ± 6.9<br>(54.5-79.5) | 77.9 ± 10<br>(66.5-92.8) | 81.4 ± 4.4<br>(74.7-89.1) | 63.3 | 64.7 ± 2.9<br>(62.0-69.6) | 72.4 ± 3.4<br>(68.8-79.7) |
| b | 4.4 ± 0.4<br>(3.8-5.4) | 4.2 ± 0.3<br>(3.8-4.9) | 9.1 ± 1.3<br>(7.8-11.7) | 11.0 ± 1.5<br>(9.5-13.5) | 8.2 ± 0.5<br>(7.7-9.3) | 8.1 ± 0.7<br>(6.7-9.3) | 8.2 ± 1.1<br>(6.5-9.4) | 8.9 ± 0.5<br>(7.7-9.8) | 6.2 | 6.1 ± 0.1<br>(5.9-6.3) | 7.7 ± 0.3<br>(7.3-8.2) |
| c | 16.9 ± 1.4<br>(15.0-19.7) | 18.2 ± 1.2<br>(17.0-19.9) | 19.4 ± 2.6<br>(16.9-26.0) | 13.0 ± 1.9<br>(10.2-16.4) | 21.6 ± 2.4<br>(18.1-25.0) | 19.4 ± 2.9<br>(15.9-26.3) | 14.9 ± 1.6<br>(12.9-17.8) | 8.3 ± 0.5<br>(7.3-11.2) | 12 | 11.4 ± 0.8<br>(10.6-12.8) | 10.2 ± 1.6<br>(8.3-13.0) |
| c' | 3.1 ± 0.1<br>(3.0-3.4) | 3.5 ± 0.2<br>(3.2-3.8) | 5.1 ± 0.6<br>(3.4-5.7) | 9.3 ± 1.8<br>(6.8-11.9) | 4.2 ± 0.4<br>(3.5-4.9) | 5.8 ± 0.6<br>(4.8-7.3) | 7 ± 0.51<br>(6.2-7.9) | 15.7 ± 1.8<br>(11.8-19) | 7.1 | 7 ± 0.6<br>(6.2-8.5) | 10.9 ± 1.9<br>(7.7-13.5) |
| T or V | 40.2 ± 6.2<br>(29.8-51.3) | 59.4 ± 1.0<br>(58.3-61.1) | 61.9 ± 9.2<br>(51.9-83.7) | 50.0 ± 1.5<br>(46.2-52.2) | 69.1 ± 6.8<br>(58.5-80.6) | 50.5 ± 1.7<br>(47.8-53.3) | - | - | 64 | 58.5 ± 5.3<br>(51.4-65.6) | 49.7 ± 2.3<br>(46.5-55.8) |
| Max. body diam. | 25 ± 4.1<br>(21-35) | 32 ± 3.4<br>(24-35) | 32 ± 8.2<br>(26-51) | 35 ± 13.0<br>(25-66) | 35 ± 5.5<br>(29-45) | 34 ± 5<br>(25-44) | - | - | 21 | 20 ± 0.9<br>(18-21) | 21 ± 0.8<br>(19-22) |
| Stoma diam. | 4.4 ± 0.6<br>(3.4-5.3) | 4.2 ± 0.6<br>(3.4-4.8) | - | - | 5.0 ± 0.8<br>(4.0-6.0) | 5.0 ± 0.9<br>(4.0-7.0) | - | - | - | - | - |
| Stoma length | 22 ± 1.2<br>(20-23) | 22 ± 1.1<br>(20-24) | 23 ± 1.4<br>(20-25) | 23 ± 2.4<br>(20-28) | 26 ± 3.4<br>(21-30) | 27 ± 5.7<br>(15-34) | 22.5 ± 1.6<br>(20-25) | 22.9 ± 1.5<br>(21-26) | 21 | 20.1 ± 0.8<br>(19-21) | 21.9 ± 1.1<br>(20-23) |
| Anterior pharynx length | 130 ± 6.8<br>(120-142) | 125 ± 7.3<br>(115-138) | 126 ± 8.0<br>(112-134) | 139 ± 24<br>(102-178) | 168 ± 28<br>(10-240) | 174 ± 17<br>(150-200) | 131.1 ± 12.8<br>(110-150) | 138.7 ± 12.1<br>(119-160) | 139 | 132 ± 10<br>(110-146) | 135.5 ± 5.4<br>(129-148) |
| Posterior pharynx length | 74 ± 4.8<br>(63-80) | 76 ± 5.1<br>(70-85) | 81 ± 7.3<br>(68-92) | 82 ± 14<br>(43-98) | 73 ± 10<br>(50-85) | 75 ± 17<br>(50-110) | 78.9 ± 7.5<br>(71-90) | 83.2 ± 4<br>(77-89) | 54 | 56 ± 6<br>(47-69) | 57.7 ± 11.1<br>(46-85) |
| Ratio of anterior and posterior pharynx | 1.8 ± 0.1<br>(1.6-2.0) | 1.7 ± 0.1<br>(1.5-1.7) | 1.6 ± 0.1<br>(1.4-1.7) | 1.7 ± 0.3<br>(1.4-2.4) | - |  | 1.7±0.1 (1.5-1.8) | 1.7 ± 0.1<br>(1.5-1.8) | 2.57 | 2.26 ± 0.3<br>(1.6-2.7) | 2.42 ± 0.4<br>(1.6-2.9) |
| Nerve ring from anterior end | 177 ± 10<br>(162-189) | 172 ± 10<br>(156-188) | 188 ± 13<br>(177-214) | 202 ± 33<br>(143-252) | - | - | - | - | 186 | 178 ± 8<br>(157-186) | 187 ± 8<br>(178-203) |
| Excretory pore from anterior end | 190 ± 12<br>(167-210) | 186 ± 13<br>(170-206) | 217 ± 19<br>(202-261) | 235 ± 49<br>(166-321) | - | 228 ± 17<br>(206-252) | 196.4 ± 14.1<br>(177-215) | 198.5 ± 19.6<br>(172-224) | 207 | 196 ± 7<br>(183-207) | 180 ± 10<br>(163-196) |
| Testis length | 408 ± 71<br>(327-561) | - | 1334 ± 409<br>(1050-2260) | - | 1493 ± 168<br>(1225-1750) | - | - | - | 851 | 746 ± 86<br>(614-866) | 362 ± 45<br>(312-423) |
| Anterior ovary length | - | 76 ± 11<br>(58-93) | - | 294 ± 174<br>(150-603) | - | 233 ± 44<br>(170-325) | - | - | - | - | 360 ± 46<br>(308-426) |
| Posterior ovary length | - | 85 ± 17<br>(65-115) | - | 268 ± 141<br>(160-560) | - | 230 ± 54<br>(130-350) | - | - | - | - | 362 ± 45<br>(312-423) |
| Anal or cloacal body diam. | 19 ± 1.8<br>(16-21) | 15 ± 1.3<br>(13.5-17.5) | 22 ± 3.3<br>(19-30) | 23 ± 6.4<br>(17-35) | 24 ± 2.8<br>(21-29) | 20 ± 2.9<br>(17-28) | - | - | 14 | 16 ± 1.8<br>(13-19) | 13.7 ± 0.4<br>(13-14) |
| Tail length | 60 ± 4.5<br>(51-65) | 53 ± 4.7<br>(46-59) | 110 ± 11<br>(98-136) | 207 ± 27<br>(163-256) | 101 ± 8.7<br>(88-116) | 118 ± 13<br>(92-138) | 118 ± 12.7<br>(95-137) | 245.8 ± 41.1<br>(177-298) | 99 | 110 ± 6<br>(99-118) | 151 ± 28<br>(108-189) |
| Spicule (curved) | 27 ± 2.4<br>(24-30) | - | 27 ± 2.1<br>(25-32) | - | 25 ± 3.6<br>(18-28) | - | 20.7 ± 2.1<br>(19-25) | - | 19 | 19 ± 1.0<br>(17-20) | - |
| Spicule (chord) | 23 ± 2.0<br>(20-26) | - | 24 ± 2.0<br>(23-29) | - | 24 ± 3.8<br>(17-30) | - | - | - | 18 | 18 ± 1.0<br>(16-19) | - |
| Gubernaculum length | 11 ± 0.7<br>(10-12) | - | 15.0 ± 1.0<br>(13-16) | - | - | - | 11.7 ± 1.3<br>(10-14) | - | 10 | 12 ± 1<br>(10-14) | - |

DESCRIPTION

### Adult

Medium-sized species with thin, slender body, strongly curved in posterior region. Cuticle striated, 0.5-1.0 µm in thickness. Lateral field with lines indiscernible. Lip region laterally flattened and expanded to form large scoop-like structure. Labial sensilla indiscernible whereas cephalic sensilla club-like. (Fig. 3.A). Stoma long and narrow *ca* 11-13 times longer than wide. Cheilostom cuticularised, long, funnel-shaped, anterior part slightly wider than posterior part. Gymnostom short, tubular, with thickened cuticularised walls, isotopic and isomorphic. Prostegostom shorter, with three fractal pieces in lateral view; mesostegostom represents considerably long part of stegostom; metastegostom slightly compressed chamber with thickened dorsal wall; metastegostomal armature comprising a short, thorn-like dorsal tooth, a medium-sized triangular right subventral tooth and thin left subventral ridge. Telostegostom sclerotised, narrow, tubular, connecting metastegostom to pharynx. Anterior part of pharynx muscular, longer than posterior part, consisting of a muscular cylindrical (procorpus) of 110-148 µm length and well developed, elongated, 46-85 µm long median bulb (metacorpus). Nerve ring surrounding posterior part of isthmus at 178-207 µm from anterior end. Excretory pore located below nerve ring at 163-207 µm from anterior end.

### Male

Body strongly arcuate in tail region. Testis single and outstretched; spermatocytes small, arranged irregularly in single or double rows at distal end. Spicule short, thick, slightly arcuate to strongly arcuate consisting of small and a reniform capitulum. Gubernaculum slender, bow-shaped to prominently keeled; weakly arcuate, tapering to blunt proximally, broad and keeled in the middle and finely attenuated at distal end. Eight pairs of genital papillae present with configuration of P1 (3.5-4 cloacal body diam. anterior to cloacal opening), P2 (1.5 cloacal body diam. anterior to cloacal opening) and P3 (just adjacent to cloacal lip), P4/P5 (located at one and a half cloacal body diam. posterior to spicules or about 1/3^rd^ of distance from spicules to tail tip, P6-7 form a group with a rudimentary bursal membrane around, P8d (located subdorsally near tail tip). Phasmids located after P8d. Tail broad and strongly arcuate ventrally. Tail tip rounded with a small mucro present terminally or subterminally.

### Female

Reproductive system didelphic, amphidelphic; anterior ovary on the right and posterior on left of the intestine. Both genital branches equally developed. Each ovary reflexed at its total length, oocytes arranged in several rows in distal half and shift to a single file proximally. Oocytes show granular texture; oviducts slender with posterior part serving as spermatheca, containing amoeboid sperms; uterus with thick walls, occasionally containing one or two egg(s). Vagina perpendicular to body axis; four small vaginal glands present, observed in lateral view; vulval lips protruding, without vulval flap. Anus a dome-shaped slit, not protuberant. Tail broad, long, weakly tapering to a rounded terminus with a small blunt mucron.

#### TYPE HABITAT AND LOCALITY

*Teratodiplogaster glomerata* n. sp. was collected from *Ficus racemosa* host trees situated in and around the campus of Indian Institute of Science, Bangalore, Karnataka, India at coordinates 13.0219° N, 77.5671° E.

#### TYPE SPECIMENS

One holotype male (*T. glomerata* n. sp./1), eight paratype males (*T. glomerata* n. sp./2-8) and nine paratype females of *T. glomerata* n. sp. on slides were deposited in Indian Institute of Science, Bangalore, Karnataka, India. One female paratype (*T. glomerata* n. sp./10) and one male paratype (*T. glomerata* n. sp./9) were deposited in the National Nematode Collection, Indian Agricultural Research Institute, New Delhi.

Etymology: The species name “*glomerata*” derived from *Ficus glomerata,* is a synonym of *Ficus racemosa*.

#### DIAGNOSIS AND RELATIONSHIP

*Teratodiplogaster glomerata* n. sp. is characterized by the presence of relatively narrow and elevated, laterally compressed lip region with fused lips, long tubular and narrow stoma with fractal pieces in prostegostom; long rectangular metacorpus; almost cylindroid to slightly expanded basal bulb; female reproductive system with conspicuous spermatheca and amoeboid sperms; males having short, arcuate spicules with tapering distal ends and curved and keeled gubernaculum; two pairs of precloacal genital papillae (P1, P2, C, P3, P4, P5d, (P6, P7), P8d, Ph) and tail conoid with a terminal or subterminal mucro. Comparison has been made with three species of *Teratodiplogaster* reported so far from *Ficus* syconia *viz., T. fignewmani* (Kanzaki *et al*., 2009) isolated form *Ficus racemosa* (Australia), *T. martini* isolated from *Ficus sycomorus* (Africa) (Kanzaki *et al*., 2012) and *T. variegatae* isolated from *Ficus variegata* (Japan) (Kanzaki *et al*., 2014).

*Teratodiplogaster glomerata* n. sp. differs from *T. fignewmani* in having smaller males (1195-1345 µm *vs* 1850-2700 µm) and females (1388-1657 µm *vs* 2160-3895 µm); females with smaller ‘b’ value (7.3-8.2 *vs* 9.5-13.5); stoma with (*vs* without) fractal dot-like arcade syncytia; amoeboid (*vs* lemon-shaped) spermatids; males having smaller ‘c’ (10.6-12.8 *vs* 16.9-26.6) and greater ‘ć’ (6.2-8.5 *vs* 3.4-5.7) values; shorter (17-20 *vs* 25-32 µm), arcuate (*vs* stout, fusiform to bow-shaped) spicules; keeled (*vs* trough-shaped) gubernaculum and two pairs [*vs* three pairs of precloacal papillae in *T. fignewmani apud* Kanzaki *et al*.(2009)].

The new species differs from *T. martini* Kanzaki *et al*., 2012 in having smaller males (1195-1345 *vs* 1825-2650 µm) and females (1388-1657 *vs* 1750-3100 µm); females with smaller ‘c’ (8.3-13.0 *vs* 15.9-26.3) and greater ‘ć’ (7.7-13.5 *vs* 4.8-7.3) values; relatively smaller metacorpus; males having smaller ‘c’ (10.6-12.8 *vs* 18.1-25.0) and greater ‘ć’ (6.2-8.5 *vs* 3.5-4.9) values; relatively shorter (17-20 *vs* 18-28 µm) spicules and keeled (*vs* L-shaped) gubernaculum and genital papillae having two precloacal pairs (*vs* three precloacal pairs) and with dissimilar configuration [P1, P2, C, P3, P4, P5d, (P6, P7) P8d, Ph *vs* P1, P2, P3, C, P4d, P5 (P6, P7), P8d, Ph in *T. martini apud* Kanzaki *et al*. (2012)].

The new species differs from *T. variegatae* Kanzaki *et al*., 2014 in having larger females (1388-1657 µm *vs* 805-1173 µm) having greater ‘a’ (68.8-79.7 *vs* 26.1-45.6), ‘b’ (7.3-8.2 *vs* 3.8-4.9) and ‘ć’ (7.7-13.5 *vs* 3.2-3.8) values; smaller ‘c’ (8.3-13.0 *vs* 17.0-19.9); lip region elevated (*vs* flattened and low); metacorpus demarcated (*vs* indistinguishable); males having greater ‘a’ (62.0-69.6 *vs* 29.9-47.4), ‘b’ (5.9-6.3 *vs* 3.8-5.4) and ‘ć’ (6.2-8.5 *vs* 3.5-4.9) values; smaller ‘c’ (10.6-12.8 *vs* 18.1-25.0); relatively shorter (17-20 µm *vs* 17-30 µm) spicules with simple, arcuate *vs* keel-like dorsal part); genital papillae with dissimilar configuration (P1, P2, C, P3, P4, P5d, (P6, P7) P8d, Ph *vs* P1, P2, P3, C, (P4, P5d), (P6, P7), P8d, Ph and tail terminus with simple [*vs* star-shaped mucro in *T. variegatae apud* Kanzaki *et al*.(2014)].

The new species differs from *T. racemosus* Bajaj & Tomar (2015) in having smaller males (1195-1345 µm *vs* 1412-2184 µm) having relatively smaller stoma (19-21 µm *vs* 20-25 µm); smaller (17-20 *vs* 19-25 µm) spicules; proximally attenuated (*vs* blunt) gubernaculum; genital papillae eight pairs (*vs* seven pairs) with dissimilar configuration (P1, P2, C, P3, P4, P5d, (P6, P7) P8d, Ph *vs* P1, P2, C, P3, P4, Ph, (P5, P6), P7d; relatively smaller females (1388-1657 µm *vs* 1587-2444 µm) with smaller ‘c’ value (10.6-12.8 *vs* 12.9-17.8); smaller (108-189 µm *vs* 177-298 µm) tail, and tail usually with subterminal (*vs* terminal mucro in *T. racemosus apud* Bajaj & Tomar, 2015).

The new species differs from *Ceratosolenus* (=*Rhabditolaimus*) *racemosa* (Anand, 2005) having presence of genital papillae (*vs* absence) and bow-shaped gubernaculum (*vs* rod-shaped). The species reported from India that include species belonging to both genus *Rhabditolaimus* and *Teratodiplogaster* have insufficient description, poor illustration and lack molecular characterization hence making it difficult to validate these as separate species. The morphometric differences within all the *Teratodiplogaster* species known so far, have been shown in Table 1.

### Molecular phylogenetic relationship

Partial SSU and D2-D3 of LSU genes were sequenced for *T. glomerata* n. sp. The relative placement of *T. glomerata* n. sp. among the other sequenced *Teratodiplogaster* and *Paradiplogaster* species was analyzed. The Bayesian tree (Fig. 8) constructed using *Koerneria luziae* (Kanzaki *et al*., 2014) as an outgroup suggested that: i) *Parasitodiplogaster* forms a monophyletic clade in relation to the *Teratodiplogaster* clade, ii) In the *Teratodiplogaster* clade, the *Teratodiplogaster* species collected from *Ficus racemosa* shows a monophyletic relation with *T. variegatae*, and iii) *T. glomerata* n. sp. forms a sister species of *T. fignewmani* and also shows a monophyletic relationship. Accession numbers in form of SSU/D2-D3 LSU for *T. glomerata* n. sp. are MT904001/ MT904002.

***Pristionchus glomerata* n. sp.**

(Figs. 9-15)

**Fig. 9.**
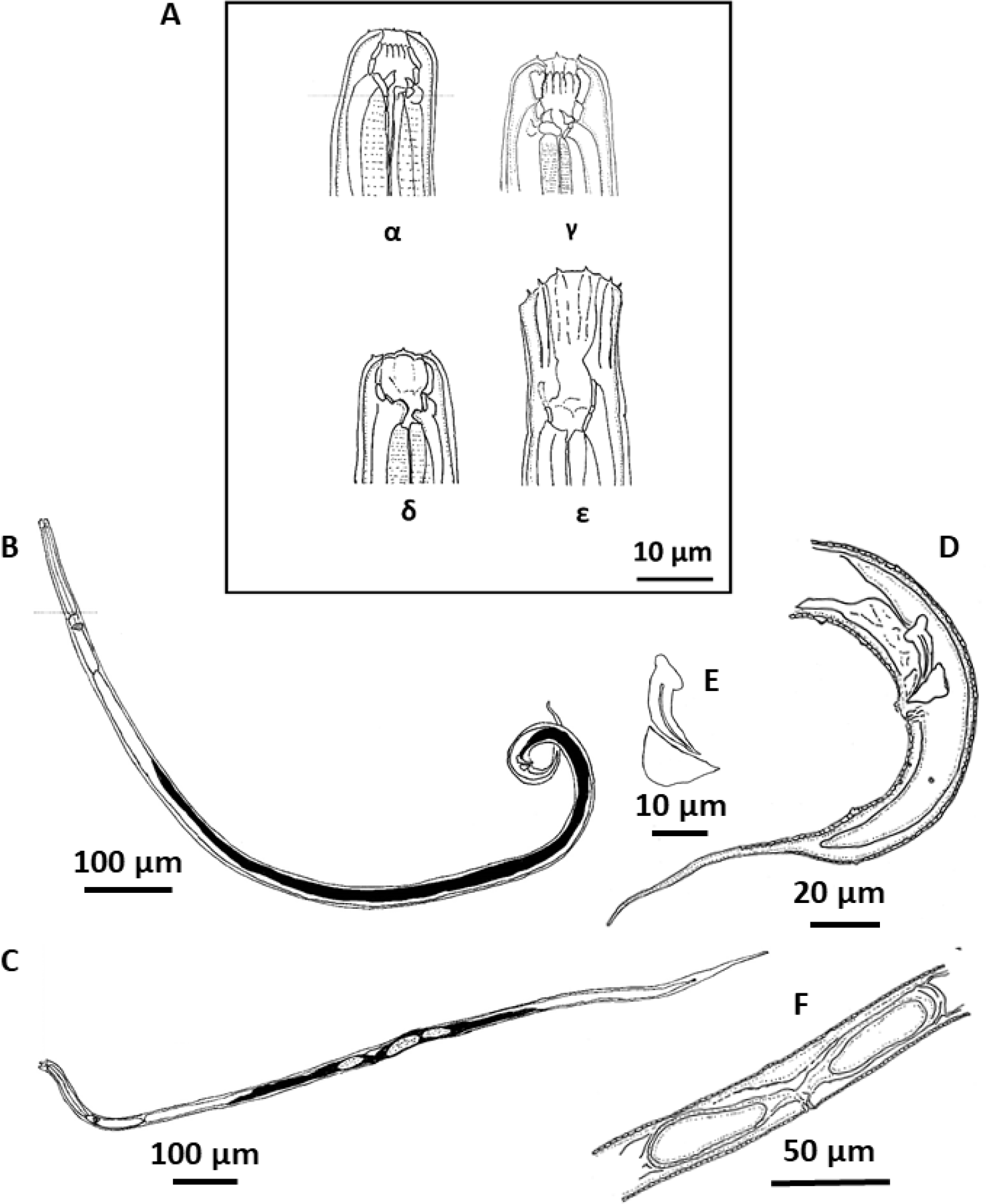
*Pritionchus glomerata* n. sp. lateral view (A) Different morphs anterior structure, (B) Habitus of male, (C) Habitus of female, (D) Tail region of male, (E) Spicule and (F) Vulval region of female.

**Fig. 10.**
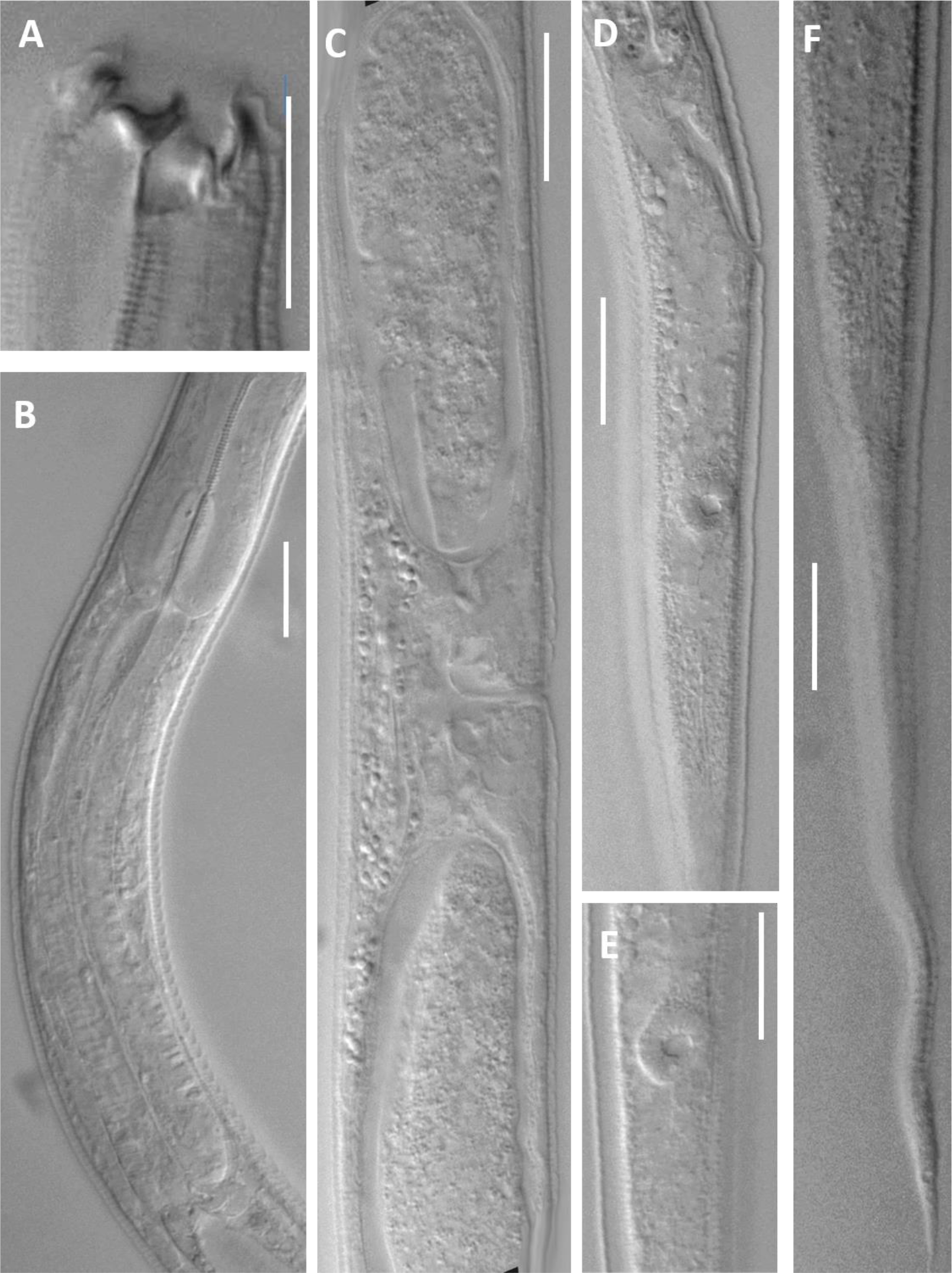
Photomicrographs of α morph (adult females only) of *Pristionchus glomerata* n. sp. (A) Head morphology, (B) Anterior and posterior pharynx, (C) different focal plane, arrows show spicule in A and capitulum in B, (C) Gonadal region of female showing vulval opening and eggs, (D) Anal opening and tail region (E – F). Scale bar: (A – F) 10 µm.

**Fig. 11.**
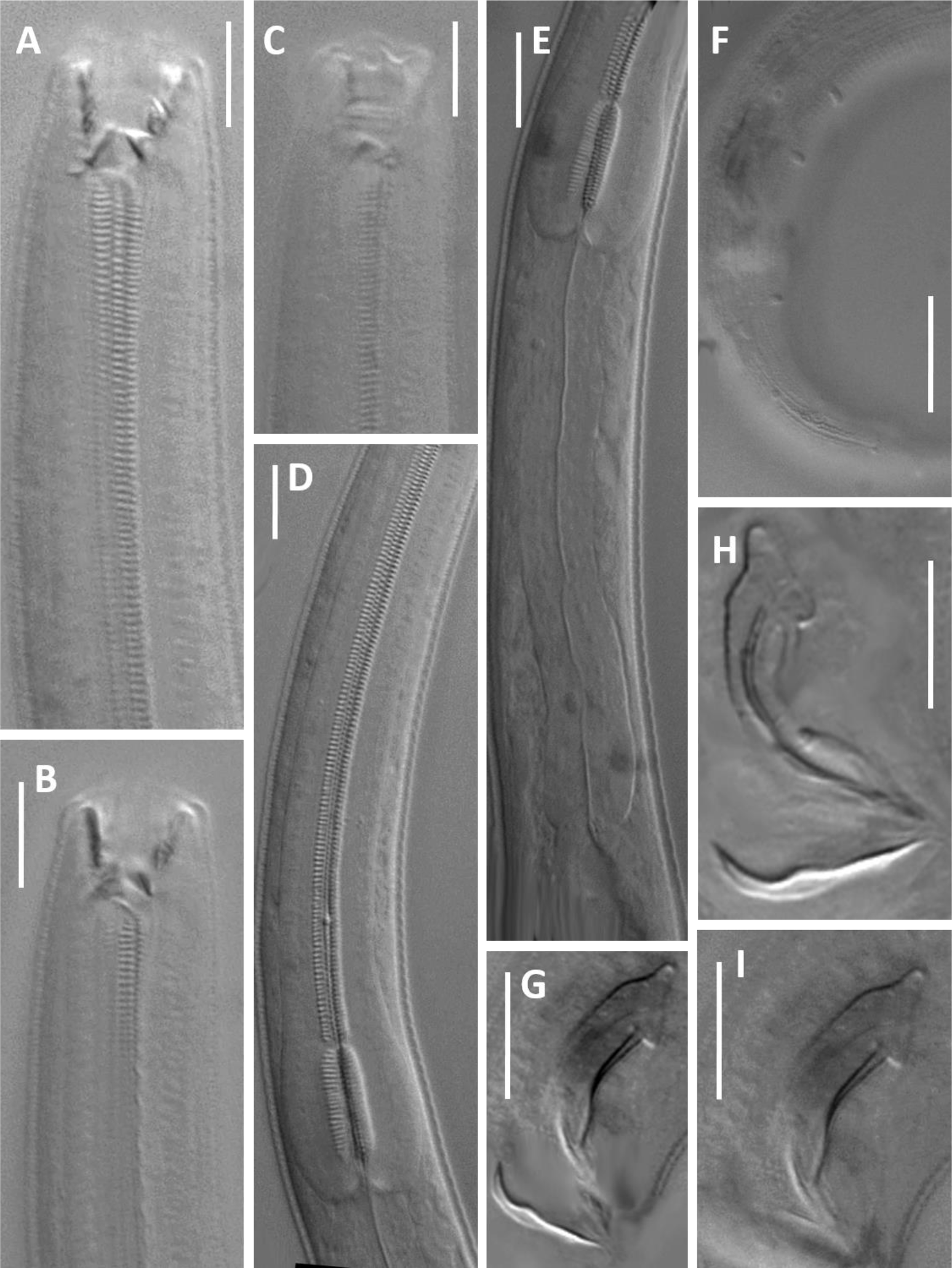
Photomicrographs of γ morph (adult males only) of *Pristionchus glomerata* n. sp. (A – C) Head morphology at different focal planes, (D) Anterior pharynx, (E) Posterior pharynx, (F) Genital papillae, (G – I) male spicule and capitulum at different focal planes. Scale bar: (A – I) 10 µm.

**Fig. 12.**
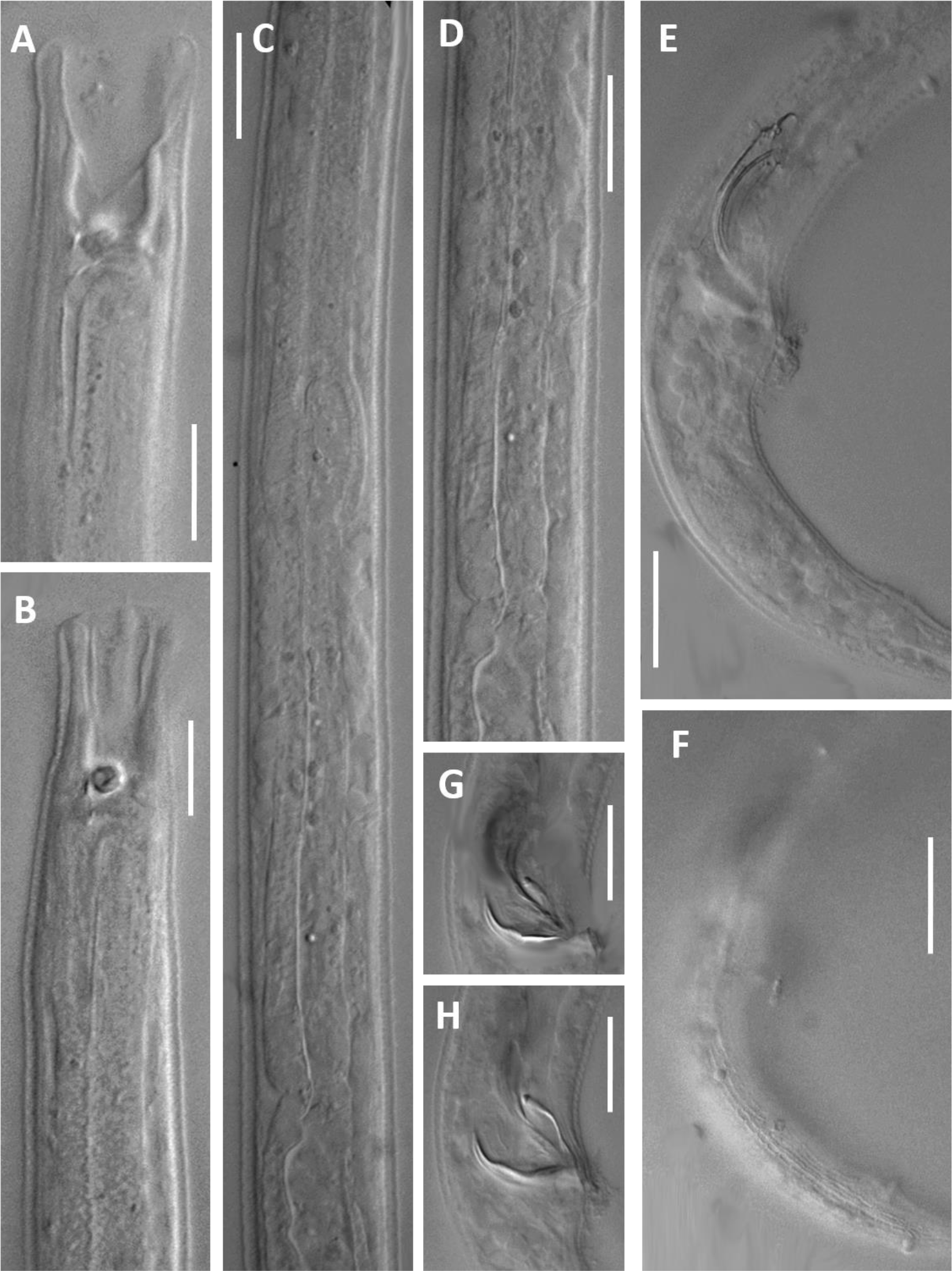
Photomicrographs of ε morph (adult males only) of *Pristionchus glomerata* n. sp. (A – B) Head morphology at different focal planes, (C – D) Anterior pharynx and posterior pharynx, (E – F) Tail region at different focal planes showing genital papillae, (G – H) male spicule and capitulum at different focal planes. Scale bar: (A – H) 10 µm.

**Fig. 13.**
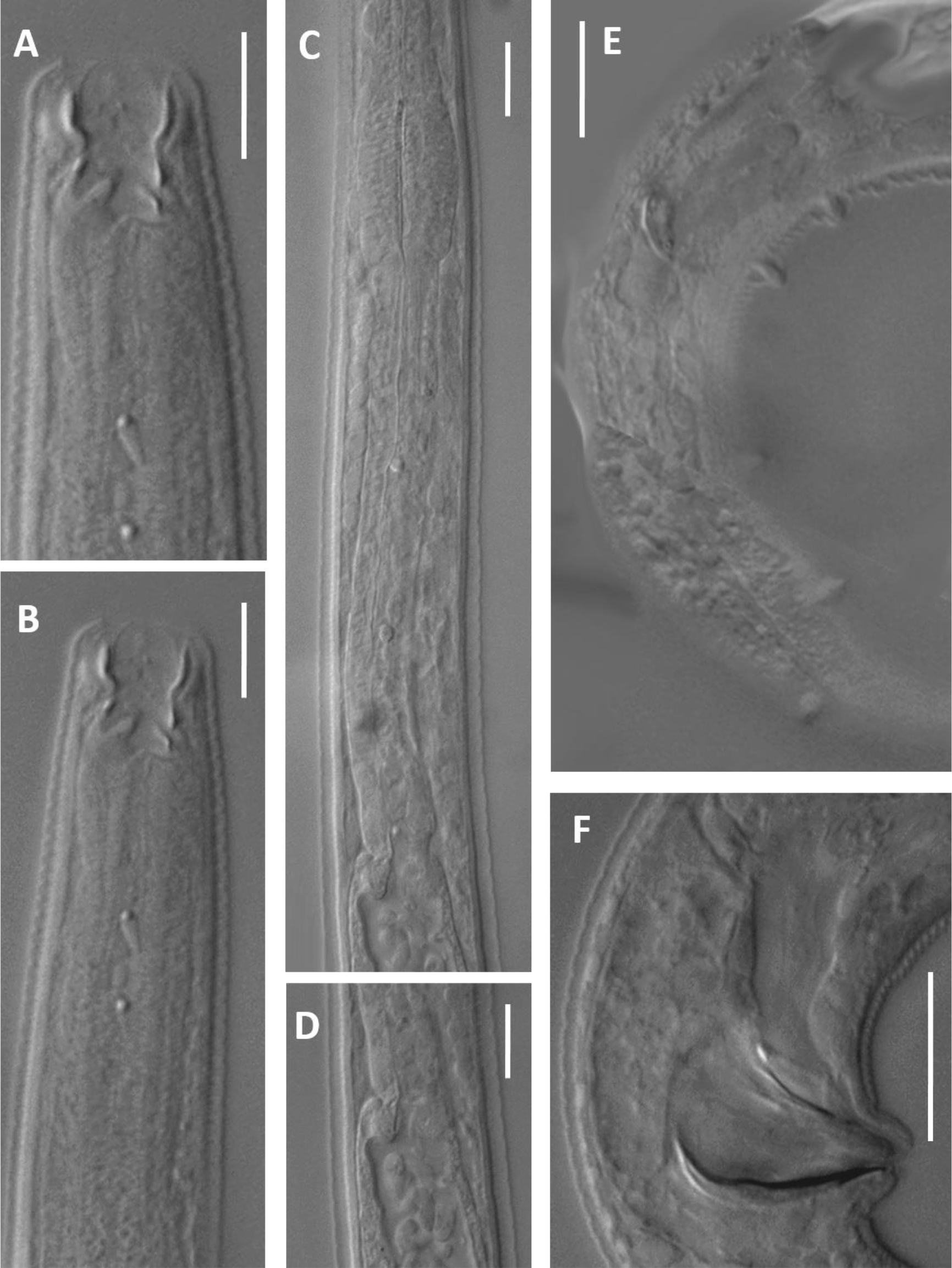
Photomicrographs of δ morph (adult males only) of *Pristionchus glomerata* n. sp. (A – B) Head morphology at different focal planes, (C – D) Anterior and posterior pharynx, (E) Genital papillae, (F) male spicule and capitulum. Scale bar: (A – F) 10 µm.

**Fig. 14.**
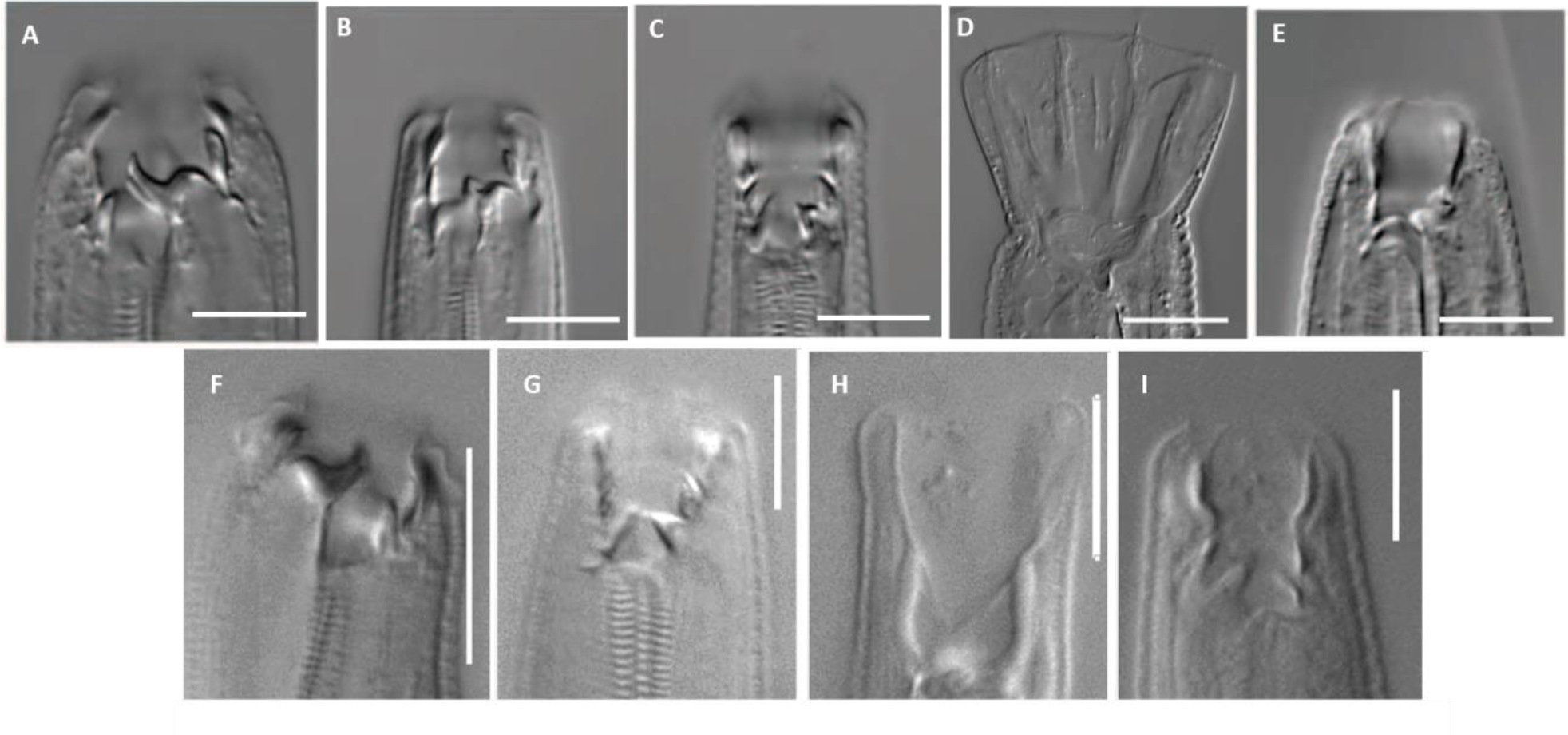
Discrete comparable adult morphs present in *P. racemosae* (A–E)[taken from Susoy *et al*. 2016) and *P. glomerata* n. sp. (F–I). The species morphs of *P. racemosae* (from left to right) : α, β, γ, ε and δ and for *P. glomerata* n. sp. (from left to right): α, γ, ε and δ Scale bar: (D) 20 µm, (A–C, E–I) 10 µm.

**Fig. 15.**
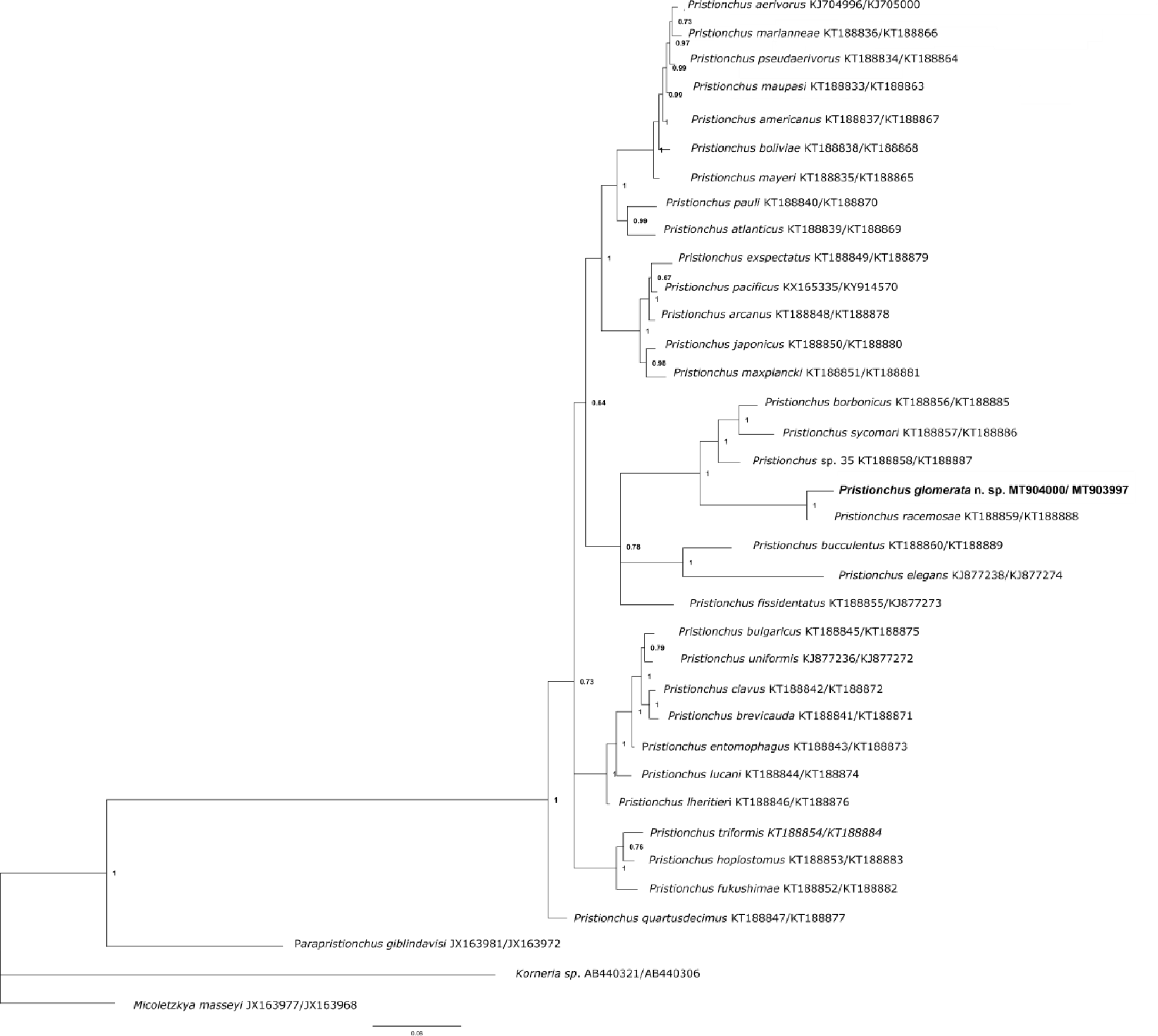
The Bayesian tree inferred from the 18S and 28S gene for *Pristionchus glomerata* n. sp. under the GTR + I + G model (lnL = 8781.2399; freqA = 0.2345; freqC = 0.1834; freqG = 0.3026; freqT = 0.2795; R (a) = 0.0543; R (b) = 0.2938; R (c) = 0.1071; R (d) = 0.0836; R (e) = 0.3843; R (f) = 0.0769; Pinva = 0.168; Shape = 0.852). The accession numbers of the compared sequences are indicated in the form: SSU/D2-D3 LSU. Posterior probability values exceeding 50% are given on appropriate clades.

MEASUREMENTS. Table 4

**Table 4.** Morphometrics of *Pristionchus glomerata* n. sp. collected from *Ficus racemosa* of Southern India. All measurements are in μm and in the form: mean ± s.d. (range).

| Character | *Pristionchus glomerata* n. sp. |  |  |
| --- | --- | --- | --- |
|  | Male |  | Female |
| | Holotype<br>($\epsilon$ morph) | Paratypes | Paratypes |
| n |  | 9 | 10 |
| L | 1253.2 | 1185.6 ± 122.6<br>(1023.8-1411.8) | 1067.7 ± 257.6<br>(693.4-1539.5) |
| a | 57.2 | 53.3 ± 6.7<br>(31.1-66.8) | 51.8 ± 10.7<br>(37.7-68.2) |
| b | 6.7 | 6.5 ± 0.6<br>(5.6-7.6) | 5.9 ± 1.2<br>(4.3-7.9) |
| c | 8.8 | 9.3 ± 1.0<br>(8.2-11.5) | 7.5 ± 0.9<br>(5.8-9) |
| c' | 7.6 | 7.1 ± 0.5<br>(6.5-7.8) | 9.2 ± 1.9<br>(7.1 -12) |
| T or V | 62.3 | 61.2 ± 4.8<br>(54-68.8) | 51.3 ± 5.6<br>(37.9-58.7) |
| Max. body diam. | 21.9 | 21.6 ± 1.5<br>(10.1-24.1) | 20.9 ± 2.9<br>(16.6-24.6) |
| Stoma length | 7.6 | 12.5 ± 6.2<br>(6.3-18.4) | 7.7 ± 2.9<br>(5.1 -14.9) |
| Anterior pharynx length | 110.9 | 109.8 ± 4.3<br>(104.2-119.4) | 101.5 ± 13.0<br>(90.3-128.2) |
| Posterior pharynx length | 74 | 73.7 ± 4.2<br>(68.1-82.3) | 75.6 ± 10.8<br>(59.2-87.1) |
| Ratio of anterior and posterior pharynx | 1.5 | 1.46 ± 0.04<br>(1.4-1.5) | 1.36 ± 0.15<br>(1.1 -1.5) |
| Neck length (head to the base of pharynx) | 192.5 | 196.0 ± 9.7<br>(185.5-217.5) | 184.85 ± 21.4<br>(158.6-230.2) |
| Excretory pore from anterior end | 53.4 | 59.4 ± 6.9<br>(52.4-75.4) | 51.9 ± 5.7<br>(46.4-63.7) |
| Anterior ovary length | - | - | 226.3 ± 44.9<br>(178.8-314.6) |
| Posterior ovary length | - | - | 268.3 ± 55.9<br>(325.1-187.2) |
| Testis length | 780.3 | 728 ± 112.9<br>(562.8-910.5) | - |
| Vulva to anus distance | - | - | 392.6 ± 170.5<br>( 783.7 -196.6) |
| Anal or cloacal body diam. | 18.7 | 17.9 ± 1.4<br>(16.1-21.4) | 15.4 ± 1.8<br>(12.2-16.8) |
| Tail length | 142.4 | 126.9 ± 9.3<br>(113.4-142.4) | 139.9 ± 23.7<br>(117.0-172.6) |
| Spicule (curved) | 21.2 | 21.42 ± 0.5<br>(20.2-22.4) | - |
| Spicule (chord) | 17.2 | 17.3 ± 0.1<br>(17.1-17.6) | - |
| Gubernaculum length | 16.5 | 16.4 ± 0.1<br>(16.2-16.8) | - |

DESCRIPTION

### Adult

Body stout, length ranging from 1-1.5 mm. Cuticle thick, transversely annulated without punctations or longitudinal striations. Lateral field showing presence of a single ridge. Stomal morphology variable in five morphotypes of the species. Anterior part of pharynx (= pro- and metacorpus) 1.5 times as long as posterior part (isthmus and basal bulb). Procorpus muscular, stout, 105.8 ± 7.3 µm long; metacorpus very muscular representing rectangular or ovoid median bulb of 43.8± 5.6 µm dimension; isthmus narrow and not muscular; basal bulb small, glandular. Pharyngo-intestinal junction conspicuous. Nerve ring encircling middle or anterior to middle region of isthmus. Excretory pore faintly visible, with position ranging from anterior level of basal bulb to pharyngo-intestinal junction. Hemizonid and deirids obscure. Five different morphs could be observed out of which the pharynx of two morphotypes (α, γ) possessed “fish-bone” or zipper-like lumen; pharyngeal lumen of morph ε is smooth. Pharynx comprising an anterior corpus continuing into a swollen metacorpus which is followed by a narrow isthmus terminating into an elongate expanded basal part.

### *Male* (general morphology comprising γ, δ, ε)

Body straight to ventrally arcuate except strongly ventrally curved posterior region. Testis single, ventrally located at 1/3^rd^ of genital branch from anterior end. Spermatocytes arranged in 2–5 rows followed by amoeboid spermatids arranged in multiple rows in genital tract. Vas deferens not clearly separated from other parts of gonad. Spicules free, bow-shaped having a wide, bilobed capitulum and pointed distal end. Gubernaculum conspicuous, flared anteriorly with ventral and dorsal walls separated at an angle of 45° with a prominent, curved dorsal wall. Tail conical, tapering down abruptly, with long spike. Nine pairs of genital papillae arranged in configuration: P1, P2d, P3, C/P4, P5, Ph (P6, P7, P8), P9d. P1 located at 1.5 cloacal body diam. anterior to cloaca, P2 was located at 1 cloacal body diam. anterior to cloaca, P3 present above the cloaca, P4 just below the cloacal opening, P5 small 4–5 µm anterior to P6 that form a group with P7 and P8, P9 posterior to P8 directed dorsally. Phasmid located between P5 and P6. Tail spike about 2.5-3.0 cloacal body diam. long.

### *Female* (morph α)

Body slightly arcuate when heat killed. Gonad didelphic, ovaries reversed. Anterior gonad right of intestine while posterior one on left side of intestine. Oocytes mostly arranged in multiple rows. Receptaculum seminis not observed; posterior part of oviduct serving as spermatheca holding small amoeboid sperm cells. Uterus spacious with 2–4 intra-uterine eggs. Vagina perpendicular to body surface; vulva elliptical slit-like with protruding lips. Posterior anal lip prolapsed. Tail long, conical to filiform. Tail spike about 2.5–3 cloacal body diam. long.

#### TYPE HABITAT AND LOCALITY

The type specimen was collected from *Ficus racemosa* host trees situated in and around the campus of Indian Institute of Science, Bangalore, Karnataka, India (GPS: (GPS: 13.0219° N, 77.5671° E).

### Morphs

Of the five morphotypes found, α morphs are represented by females while γ, δ, ε are represented by males. The morphs mainly show phenoplastic traits in the anterior region especially the labial region, stoma and pharyngeal lumen. However, the general characters of females remain the same while males also show similar spicules and gubernaculum morphologies. Morphological illustrations and photographs are shown in Figs. 9 (A), 10-15.

Morph α (Females): Lip region wide, slightly offset with six equal-sized, rounded lips; cheilostom equal or slightly longer than gymnostom, absence of tightly packed cheilostomal rugae, cheilostomal flap weak; gymnostom with prominent serrated anterior margin; pro-mesostegostom without lobes; metastegostom with claw-like dorsal tooth, curved right subventral tooth, and coarsely serrated left subventral ridge overlapping 2/3^rd^ of the dorsal tooth; telostegostom long, sclerotized with straight dorsal and wide tapering subventral walls. Pharyngeal lumen zipper-like. Conspicuously larger phasmids present in anterior half of the tail, not found in any other morph.

Morph γ (Male): Lip region narrow; lips distinct; six equal-sized, amalgamated, stoma funnel-shaped, wider anteriorly and tapering at base; cheilostom thick; gymnostom relatively thin, smooth, narrower; metastegostom with a large flap-like dorsal tooth, a triangular right subventral tooth and left subventral ridge; stegostom simple, smooth, continuing into zipper-like (fish bone-like) pharyngeal lumen.

Morph δ (Male): Lip region with six equal-sized, amalgamated lips; cheilostom thick with sloping walls, arched; gymnostom half the length of the cheliostom, thickened, arched; metastegostom with a triangular dorsal tooth and right subventral, dagger-like tooth and a left subventral plate; telostegostom shallow, short, weakly sclerotized.

Morph ε (Male): Anterior labial region with umbrella-like flap having six labial projection; left side possessing a slit with a round opening at base, the flap in few specimens covers the gymnostom and cheilostom of one side and with six ribs consisting of labial papillae within the cuticle; cheilostom smooth, anterior margin marked by per- and interradial notches; cheilostom and gymnostom equal in length and project outward, metastegostom simple, smooth, except for a flat, thin, claw-like dorsal tooth, ventral tooth absent; telostegostom weakly sclerotized, indistinct.

#### TYPE HABITAT AND LOCALITY

*Pristionchus glomerata* n. sp. was collected from *Ficus racemosa* host trees situated in and around the campus of Indian Institute of Science, Bangalore, Karnataka, India at coordinates 13.0219° N, 77.5671° E.

#### TYPE SPECIMENS

One holotype male ε morph (*P. glomerata* n. sp./1), eight paratype males (*P. glomerata* n. sp./2-8) and nine paratype females (*P. glomerata* n. sp./1-9) of *P. glomerata* n. sp. on slides were deposited in Indian Institute of Science, Bangalore, Karnataka, India. Holotype male (ε morphotype) (*P. glomerata* n. sp./9) was deposited in the National Nematode Collection, Indian Agricultural Research Institute, New Delhi.

Etymology: The species name “*glomerata*” derived from *Ficus glomerata,* is a synonym of *Ficus racemosa*.

#### DIAGNOSIS AND RELATIONSHIP

*Pristionchus glomerata* n. sp. is characterized by its phoretic relationship with *Ceratosolen fusciceps*, the pollinator wasp of the *Ficus racemosa,* and presented four different morphotypes *viz*., α, γ, δ and ε types. The morphs mainly show variations in labial region ranging from rounded, truncate to umbrella-shaped, stoma from barrel-shaped to cuboidal or elongated, dorsal tooth being claw-shaped to triangular, dagger-like and pharyngeal lumen simple to zipper-like; female reproductive system didelphic, ovaries reversed; oocytes mostly arranged in multiple rows; proximally dilated oviduct holding amoeboid sperm cells; 2–4 intra-uterine eggs occasionally present; vulva elliptical slit-like with protruding lips; males having a bow-shaped spicules with a wide, bilobed capitulum; gubernaculum conspicuous, with 45° bent, curved dorsal wall. Tail conical, tapering down abruptly, with long spike. Nine pairs of genital papillae arranged in configuration P1, P2/ P2d, P3, C/P4, P5, Ph, (P6, P7, P8), P9d. The morphotype α possessed exceptionally large phasmids.

The new species is distinguished from all other Diplogastridae, except *P. borbonicus* Susoy *et al*., 2016, *P. sycomori* Susoy *et al*., 2016 and *P. racemosae* Susoy *et al*., 2016 by the presence of morphs of laterally symmetrical and laterally asymmetrical stomatal structures. The diagnostic characteristic features of this species are the gross morphological differences in the morphs compared with those of the species reported from fig species. Among the fig-associated Pristionchus species, P. glomerata n. sp. comes close to P. racemosae in having the basic structural similarity in the stomal components of four morphs; however, relatively cylindrical (vs globular) buccal cavity in γ and δ morphotypes and relatively rectangular and less expanded (funnel-shaped and well expanded) lip region in ε morphotype in *P. racemosa*e and lacks β morph.

*P. glomerata* n. sp. differs from *P. sycomori* in having lips moderately developed (*vs* large), equal-sized (*vs* lateral lips higher and wider than subventral and subdorsal lips); labial region with ring of conspicuous, thin cuticular cheilostomal filaments); cheilostom (equal or smaller *vs* twice the length of gymnostom) with less developed cheilostomal rugae (*vs* thick, tightly packed rugae); gymnostom without (*vs* with) heavy punctations, lip region slightly narrower (*vs* conspicuously wider) than the adjoining body; anterior labial margin smooth (*vs* projected, “wave”-like) in α morphotype and larger, relatively expanded and umbrella-like (*vs* less expanded, barrel-shaped) lip region in ε morphotype in *P. sycomori apud* Susoy *et al*. (2016) (Fig. 15).

*Pristionchus glomerata* n. sp. differs from *P. borbonicus* in having lips moderately developed (*vs* large), equal-sized (*vs* lateral lips higher and wider than subventral and subdorsal lips); labial region with ring of conspicuous, thin cuticular cheilostomal filaments); cheilostom (equal or smaller *vs* twice the length of gymnostom) with less developed cheilostomal rugae (*vs* thick, tightly packed rugae); gymnostom without (*vs* with) heavy punctations, lip region slightly narrower (*vs* conspicuously wider) than the adjoining body; anterior labial margin smooth (*vs* projected, “wave”-like) in α morphotype and larger, relatively expanded and umbrella-like (*vs* less expanded, barrel-shaped) lip region in ε morph in *P. sycomori apud* Susoy *et al*. (2016). *Pristionchus glomerata* n. sp. differs from *Canalodiplogaster racemosus* (Bajaj & Tomar 2015) in having 9 pairs of genital papillae (*vs* 7 pairs), conspicuous gubernaculum (*vs* large pouch shaped) and spicule monomorphic (*vs* dimorphic).

#### REMARKS

Bajaj and Tomar (2015) reported a new genus *Canalodiplogaster* with *C*. *racemosus* as its species from the fig *F. racemosa* along with three other new genera. The description and illustrations of the former indicated it to be a species of *Pristionchus* with three morphotypes. The authors have identified the morphs as stenostomous and eurystomous individuals. The present species *P. glomerata* n. sp. with five morphs can be distinctly differentiated from the species (= *C. racemosus*) in having γ morph with funnel-shaped (*vs* barrel-shaped) buccal cavity; smaller (20-22 µm *vs* 29-43 µm) spicules and morphotype ε with umbrella-like lip region represented by only males (*vs* females in the species (=*C. racemosus*) *apud* Bajaj and Tomar (2015). The status of the latter species can be verified by revisiting the species for the molecular data and for detailed information about all the morphotypes associated with the species.

### Molecular phylogenetic relationship

Partial SSU and D2-D3 of LSU genes were sequenced for *P. glomerata* n. sp. The relative placement of *P. glomerata* n. sp. among the other known sequenced *Pristionchus* species was analyzed. The Bayesian tree (Fig. 16) constructed using *Koerneria* sp. (Kanzaki *et al*., 2014), *Parapristionchus giblindavisi and Micoletzkya masseyi* (Susoy *et al*., 2016) as an outgroup, suggested that: i) The fig-associated *Pristionchus* form a polyphyletic clade in relation to *P. bucculentus, P. elegans* and *P. fissidentatus*, ii) In the fig-associated *Pristionchus* clade, *Pristionchus* collected from *Ficus racemosa* shows a monophyletic relation to other fig-associated *Pristionchus* species, and iii) *P. glomerata* n. sp. is a sister species of *P. racemosae* and also shows a monophyletic relationship. Accession numbers in form of SSU/D2-D3 LSU for *Pristionchus glomerata* n. sp. are MT904000/ MT903997.

## Discussion

Nematodes have long been associated with the fig syconium as a substratum for their growth and reproduction whereas the pollinators associated with figs are used as vehicles for their transport from one syconium to the other (Krishnan *et al*., 2010). These nematodes in association with the fig syconium have shown millions of years of co-evolution, co-speciation and co-diversification (Herre, 1993; Davies *et al*., 2015). Such associations might range from being commensal to parasitic in nature (Giblin-Davis *et al*., 2013). The fig-associated nematodes of families Aphelenchoididae and Diplogastridae have shown independent phylogenetic radiation (Kanzaki *et al*., 2009; Davies *et al*., 2015). The former family constitutes plant-parasitic nematodes that feed on the anthers and the epidermis of the female florets of the fig (DeCrappeo & Giblin-Davis, 2001; Vovlas *et al*., 1992; 1996; 1998; Giblin-Davis *et al*., 1995, 2006; Center *et al*., 1999) whereas the latter family includes nematodes which can be fungal feeders, bacteriovores or insect parasites (Susoy *et al*., 2016).

*Ficus racemosa*, the fig plant under study in this paper, is a monoecious species of subgenus *Sycomorus* commonly found in Bangalore, South India. A single pollinator wasp species *Ceratosolen fusciceps* and six non-pollinator fig wasp species (Ghara & Borges, 2010) have been found associated with this fig species, of which the former serves a reliable vehicle for nematode transport (Krishnan *et al*., 2010, Gupta & Borges, 2019). So far, the nematode species reported from *Ficus racemosa* belong to genera *Schistonchus, Ficophagus*, *Teratodiplogaster and Pristionchus* (*=Canalodiplogaster*) (Reddy & Rao, 1986; Anand, 2002; Anand, 2005; Bajaj & Tomar, 2014, 2015). However, in the present study we could find only three species associated with *Ficus racemosa*, each a representative of *Ficophagus*, *Teratodiplogaster* and *Pristionchus.* Coincidentally, all the species described from India lack molecular characterization and are differentiated mainly on minor morphological characters without any report of phenotypic plasticity thus causing difficulty in ascertaining their correct status in comparison with other species (Anand, 2002; Anand, 2005; Reddy & Rao, 1986; Davies *et al.,* 2013; 2015). *Ficophagus glomerata* n. sp. forms a sister species with *F. microcarpus* isolated from *Ficus microcarpa* in China under the phylogenetic analysis done using partial SSU region and *Ficophagus* sp. isolated from *Ficus obliqua* using the D2–D3 segment of LSU. It was found to be distantly related to the species found in Australia suggesting that the species might have diverged out separately either because of geographical range variation or due to host shift. The phylogenetic trees stand in concordance with the earlier study where it was proposed that the genera *Schistonchus and Ficophagus* represented two evolutionary lines of the polyphyletic clade (Davies *et al*., 2010).

The diplogastrid species, *T. glomerata* n. sp. shows affinities with *T. fignewmani* and *T. racemosus*. The presence of *T. glomerata* n. sp. in *Ficus racemosa* syconia suggests that a speciation event might have occurred due to large geographical ranges and genetic isolation of *F. racemosa* in south India compared to the south-east Asian populations (Bain *et al.,* 2016). The phylogenetic analysis done using SSU and LSU shows that *Teratodiplogaster* forms a monophyletic clade with respect to the *Pristionchus* clade. The new species *T. glomerata* n. sp. forms a sister species to *T. fignewmani*. The data still stands in concordance with the earlier study suggesting the *Parasitodiplogaster* Poinar, 1979, and *Teratodiplogaster* Kanzaki *et al*. 2009 clades to be monophyletic and separated into five groups, i.e. *Parasitodiplogaster australis* Bartholomaeus, *et al*. 2009, *Parasitodiplogaster sycophilon* Poinar 1979, *Parasitodiplogaster maxinema* Poinar & Herre, 1991, *Parasitodiplogaster laevigata* Giblin-Davis *et al*. 2006 and *Parasitodiplogaster citrinema* Poinar & Herre, 1991, and the *Teratodiplogaster* Kanzaki *et al*. 2009 clade (Wöhr *et al*., 2014).

*Pristionchus*, another genus of Diplogastridae, is known to be mainly associated with beetles but recently has been reported from *Ficus* species (Herrmann *et al*., 2006, 2007; Kanzaki *et al.,* 2011, 2012a, 2012b). The species found so far are *P. borbonicus* collected at Grand Étang, La Réunion Island, from *F. mauritiana*; *P. sycomori* collected in South Africa from *F. sycomorus*; *P. racemosae* collected in Vietnam from *F. racemosa* (Susoy *et al*., 2016) and a species of *Pristionchus* reported (Bajaj & Tomar, 2015) from India with the name *Canalodiplogaster racemosus*. According to the phylogenetic analysis, *Pristionchus glomerata* n. sp. forms a closely related sister species to *P. racemosae* where the branch length increase in the arm of *Pristionchus glomerata* n. sp. depicts speciation with recent divergence. The species present in *Ficus racemosa* forms a monophyletic clade to the other *Pristionchus* species which are associated with figs. The close association is suggestive of speciation which might have occurred due to large geographical range variations. The occurrence of five morphotypes is an ideal case of character displacement for the efficient sharing of resources in the microhabitat of syconium populated by a good number of species. The morphs (δ, ε) with simple, tube-like buccal cavities devoid of effective armature seem to compete for a temporary bacterial food source whereas the predatory morphs (α, γ) emerge next coinciding with the proliferation of wasp-transmitted nematodes in the syconium. Development of greater number of morphs might be due to the high climatic and seasonal variation which is observed in southern India (Gadgil & Joshi, 1983; Gunnel 1997; Peel *et al*., 2007; Shenoy & Borges, 2010; Chanam *et al.,* 2014). We observed sex specific stomatal morphs (β = adult female only) versus (γ,δ, ε = adult males only) which requires further validation through a study similar to that reported in Susoy *et al*. (2016) with *Pristionchus sycomori* where different early and late interfloral (phase C) figs were examined and large numbers of *Pristionchus* were sexed and assessed for stomatal morphotype. This will be attempted in the future.

The phylogenetic analyses of these nematodes using LSU and SSU of rRNA sequences have helped in the better understanding of nematode diversification. There are instances in which a single *Ficus* species is known to be associated with several nematode species belonging to the same genus, for example, several species of *Schistonchus* associated with *Ficus racemosa* in Australia (Davies *et al*., 2010), which is suggestive of opportunities for host switching for such cosmopolitan fig species (Zhang *et al*., 2006). Future studies on nematode taxonomy related to *Ficus* species should involve sequencing of at least two genes which might help to better resolve the phylogenetic trees of these nematode species. Special emphasis should be given to comparative genomic analyses using the new species and its close relatives which would yield interesting information concerning the genes involved in different life history traits, switching feeding habitats and the development of plant and animal parasitism. Further investigation on *Pristionchus* species associated with *Ficus* species might help us to conduct polymorphism studies across the several geographical landscapes to understand the relationships between form and function and how evolution may have proceeded from a common ancestor. This might even lead to better understanding of life history adaptations used by these nematode species. This study also paves the way for further research on fig nematodes related to their biodiversity, distribution, evolutionary history, host/carrier relationships with pollinators associated with the *Ficus* systems and co-speciation between fig and fig associated nematodes.

## Acknowledgements

This research was supported by funds from the Department of Biotechnology-Indian Institute of Science (IISc) Partnership Programme, and the Ministry of Environment, Forests & Climate Change. We are grateful to G Yettiraj and Sunitha Murray for logistical support. We are grateful to the University Sophisticated Instruments Facility (USIF) at Aligarh Muslim University for providing the electron microscope images. We thank the Bio-Imaging Facility at the Biological Sciences Division, IISc, for confocal images. We are grateful to two anonymous reviewers for their critical comments.

